# Multiscale PHATE Exploration of SARS-CoV-2 Data Reveals Multimodal Signatures of Disease

**DOI:** 10.1101/2020.11.15.383661

**Authors:** Manik Kuchroo, Jessie Huang, Patrick Wong, Jean-Christophe Grenier, Dennis Shung, Alexander Tong, Carolina Lucas, Jon Klein, Daniel Burkhardt, Scott Gigante, Abhinav Godavarthi, Benjamin Israelow, Tianyang Mao, Ji Eun Oh, Julio Silva, Takehiro Takahashi, Camila D. Odio, Arnau Casanovas-Massana, John Fournier, Yale IMPACT Team, Shelli Farhadian, Charles S. Dela Cruz, Albert I. Ko, F. Perry Wilson, Julie Hussin, Guy Wolf, Akiko Iwasaki, Smita Krishnaswamy

**Affiliations:** Department of Neuroscience, Yale University, New Haven, CT; Department of Computer Science, Yale University, New Haven, CT; Department of Immunobiology, Yale University, New Haven, CT; Montreal Heart Institute, Montréal, Québec, Canada; Department of Medicine, Yale University, New Haven, CT; Department of Genetics, Yale University, New Haven, CT; Computational Biology, Bioinformatics Program, Yale University, New Haven, CT; Department of Applied Mathematics, Yale University, New Haven, CT; Department of Epidemiology of Microbial Diseases, Yale School of Public Health, New Haven, CT; Department of Medicine, Section of Infectious Diseases, Yale University School of Medicine, New Haven, CT; A list of authors and their affiliations appears at the end of the paper; Department of Medicine, Section of Pulmonary and Critical Care Medicine, Yale University School of Medicine, New Haven, CT; Clinical and Translational Research Accelerator, Department of Medicine, Yale University, New Haven CT; Faculty of Medicine, Université de Montréal, Québec, Canada; Mila – Quebec AI institute, Montréal, Quebec, Canada; Department of Mathematics and Statistics, Université de Montréal, Montréal, Quebec, Canada; Howard Hughes Medical Institute, Chevy Chase, MD, USA

**Author notes:** Equal contribution (Manik Kuchroo, Jessie Huang, Patrick Wong). Jointly supervised work (Julie Hussin, Guy Wolf, Akiko Iwasaki, Smita Krishnaswamy). Correspondence to Smita Krishnaswamy, 333 Cedar St, New Haven, CT 06520. E-mail:; and to Akiko Iwasaki, 300 Cedar St, New Haven, CT 06520.

**Keywords:** manifold learning, machine learning, visualization, clustering, COVID-19, single cell

## Abstract

The biomedical community is producing increasingly high dimensional datasets, integrated from hundreds of patient samples, which current computational techniques struggle to explore. To uncover biological meaning from these complex datasets, we present an approach called Multiscale PHATE, which learns abstracted biological features from data that can be directly predictive of disease. Built on a continuous coarse graining process called diffusion condensation, Multiscale PHATE creates a tree of data granularities that can be cut at coarse levels for high level summarizations of data, as well as at fine levels for detailed representations on subsets. We apply Multiscale PHATE to study the immune response to COVID-19 in 54 million cells from 168 hospitalized patients. Through our analysis of patient samples, we identify CD16^*hi*^ CD66b^*lo*^ neutrophil and IFNγ^+^GranzymeB^+^ Th17 cell responses enriched in patients who die. Further, we show that population groupings Multiscale PHATE discovers can be directly fed into a classifier to predict disease outcome. We also use Multiscale PHATE-derived features to construct two different manifolds of patients, one from abstracted flow cytometry features and another directly on patient clinical features, both associating immune subsets and clinical markers with outcome.

## 3 Introduction

Extremely high throughput biomedical data is generated by a range of technologies [1–6] that measure dozens to tens of thousands of features in millions of individual cells. Furthermore, these technologies are now applied to large patient cohorts, providing information that must be integrated and analyzed at scale to provide insights into cellular mechanisms and patient responses. However, there are no specific methods designed to sift through such data at varying levels of granularity to uncover features that are directly associated with disease phenotype. The SARS-CoV-2 pandemic has brought this problem to the forefront of biologists’ minds. As increasingly large datasets are built by integrating patient samples from around the globe, computational approaches also must scale to provide improved insights regardless of technology type.

We posit here that the key to understanding such vast and complex data is to create meaningful representations that uncover structure at all resolutions or scales. This approach involves learning representations of the biological system at many levels, allowing for coarse, high level summarization as well as fine grained, detailed representations of data subsets. Current tools for dimensionality reduction and data exploration - including *t*-distributed stochastic neighborhood embedding (tSNE) [7], uniform manifold approximation and projection (UMAP) [8], as well as principle component analysis (PCA) [9] - only show a single level of granularity of the data. Recent computational papers on SARS-CoV-2 have represented data using one of these approaches [10, 11], visualizing the major cell types such as B cells, T cells and myeloid cells. Differences between an effective immunological response and an ineffective one, however, may not be found at the granularity of immune compartment abundance alone. In fact, appreciation of a finer resolution of the T cell manifold would reveal subsets that may be predictive of disease severity. This phenomenon is found across biomedical data science, as the state space of the data is generally a collection of manifolds or continuum structures which can be organized at varying levels of hierarchy.

Based on this insight, we developed Multiscale PHATE, a method that can learn and visualize abstract cellular features and groupings of the data at *all levels of granularity*. Our algorithm is based on a dynamic process we have developed called diffusion condensation [12], which computes a manifold-intrinsic diffusion space on the original data before slowly condensing data points towards local centers of gravity to form natural, data-driven groupings across multiple granularities. Rather than forcing merges at each iteration, as done in most agglomerative hierarchical clustering methods, diffusion condensation allows cells to naturally come together over the course of successive condensation steps before merging points that fall within a threshold. Visualizing a series of iterations in this dynamic condensation process using the manifold-affinity preserving PHATE method creates Multiscale PHATE embeddings, while evaluating groupings of cells across granularities naturally creates Multiscale PHATE clusters. Furthermore through efficient scalable implementation, we show that we are able to perform condensation, visualization and clustering of large-scale the data significantly faster than “single-scale” visualization techniques like tSNE, UMAP or our earlier method PHATE [13].

We showcase our method on 251 blood samples from 168 patients infected with SARS-CoV-2 measured across four different flow cytometry panels. Analysis of 54 million of cells by single resolution dimensionality reduction and clustering algorithms would take days to weeks to perform. With our unique multigranular approach, we can produce high level summarizations as well as detailed cell type specific analyses of each panel of markers within minutes. When combined with the MELD [14], we find that our approach is particularly powerful at identifying cannonical and non-cannonical cellular populations associated with patient outcome across resolutions. At coarse resolution, we identify T cells to be broadly protective while monocytes and granulocytes to be pathogenic. At finer resolution, we identify unique non-cannonical populations of cells, such as D16^*hi*^ CD66b^−^ neutrophil, CD14^−^CD16^*hi*^ HLA-DR^lo^ monocytes, and IFNγ^+^GranzymeB^+^ Th17 cells, to be associated with patient mortality. This type of multigranaular analysis reveals that though broadly a cell type, such as T cells, may be protective, fine grain analysis reveals cellular subsets that can be pathogenic, highlighting the need for a multiresolution approach. Next, we show that Multiscale PHATE-derived cellular groupings can be used as features input to a random forest classifier to predict outcome better than immunologist curated and gated features. A unique contribution we make is the use of these multiscale feature proportions as descriptors of each patient, which can be used to create a patient-level embedding. We relate distances between these feature descriptors to Earth Mover’s Distance between patients, leading to an robust patient embedding.

Finally, to display the generality of our approach across data types, we created a multiscale distillation of clinical data from 2,135 patients admitted to Yale New Haven Hospital (YNHH). Built from 18 laboratory, clinical, and demographic variables, Multiscale PHATE was able to create multiresolution embeddings of patient clinical states and identify regions enriched for different patient outcomes. By associating clinical features and cellular populations with outcomes, we found markers of multi-organ dysfunction to be associated with mortality and overall T cell counts to be associated with length of recovery from infection.

## 4 Results

### 4.1 Multiscale PHATE Algorithm

Multiscale PHATE is a multiresolution visualization and clustering method that can reveal the structure of data visually and quantitatively at multiple granularities. Multiscale PHATE adapts and combines two powerful algorithms, diffusion condensation for multigranular analysis and PHATE for dimensionality reduction, to produce visualizations and clusters at all levels of granularity.

Diffusion condensation is a dynamic process by which data points slowly and iteratively come together at a rate determined by the diffusion probabilities between them [12]. This iterative process is powerful as it reveals structure and groupings of the data at all levels of granularity for a given diffusion kernel bandwidth. The diffusion condensation process involves three steps that are iteratively repeated until all points converge:

1. Compute a Markov diffusion operator from the data
2. Apply this operator to the data as a low pass filter, moving points towards local centers of gravity
3. Merge points together that fall below a preset distance threshold

The first step involves computing a Markov diffusion operator from a dataset. This is done by first computing a distance matrix **D** between all data points, before converting this to an affinity matrix **K** by using an fixed bandwidth Gaussian kernel function as done previously [12]. **K** is then row normalized to obtain a *diffusion operator* between data points. In the next step, the diffusion operator is applied to the input data (the data matrix is left-multiplied by the diffusion operator), effectively replacing the value of a point, with the weighted average of its diffusion neighbors. This causes points to *condense*, or move closer together due to the removal of variation. In graph spectral terms this step is akin to applying a *low-pass filter* on the graph frequency spectrum which smooths the data. In the third step, diffusion condensation merges points that have condensed within a preset merge threshold together. This process is then repeated iteratively. At each step, diffusion operator is calculated on the output of the previous iteration and used as a low-pass filter to produce an increasingly coarse dataset for the next iteration. Over the course of many iterations, data points slowly converge to local centers of gravity and collapse into each other, effectively grouping them together at that level of granularity. As the diffusion condensation process continues, we scale the diffusion operator to diffuse points to more global centers of gravity. This first removes local variability in the data at initial iterations before removing more global variability in later iterations. This deep cascade of low pass filters effectively builds a tree by merging data points together in a natural manner.

In its original form, the diffusion condensation process does not scale to millions of data points, does not condense points on a manifold, and is not optimized for visualization. Thus, we have modified diffusion condensation to allow for scalable and effective visualizations. The main steps of Multiscale PHATE include:

1. Transforming datapoints to a novel diffusion potential coordinate system to learn the data manifold,
2. Computing a *fast* diffusion condensation process that scales to millions of data points,
3. Identifying levels for visualization based on gradient analysis and creating a density aware visualization.

Each of these steps is explained below.

#### The diffusion potential coordinate system

The original diffusion condensation algorithm [17] implements the coarse graining of the data in the ambient measurement space. However, condensing in this space can lead to “averaged” points that deviate from the intrinsic data manifold, especially in cases where the intrinsic manifold is very curved (Supplementary Figure 1A). As cellular state spaces can be heavily non-linear [13, 15, 18], we require an alternative method of diffusion condensation that ensures that the condensed points remain on the manifold. A straightforward method for achieving this might be diffusion map coordinates. However, the computation of diffusion map coordinates requires eigendecomposition of a diffusion operator which is slow (*O*(*n*^3^) complexity).

As we do not need to reduce dimensions, we could simply run diffusion condensation on the diffusion operator of the dataset itself. However, as discovered in [13], diffusion distances have a key drawback: they are dominated primarily by differences in nearest neighbors and ignore more distant datapoints. For example, when visualizing two cells that share nearest neighbors, distant cells that could be very useful in their global placement are ignored. This was remedied in [13] by computing the *diffusion potential*, which *log* scales diffusion probabilities to allow for faraway datapoints to inform the local distances and impact the global geometry of the visualization. Inspired by this finding, we chose diffusion potential as a coordinate system for cells in our multiscale approach. These coordinates are a way of re-representing each cell in the dataset by the potential of the diffusion probabilities to other datapoints and offers a “straightened” and globally coherent intrinsic manifold space upon which to operate the diffusion condensation process.

#### Fast diffusion condensation

To achieve scalability, we employ two techniques to improve computational efficiency. First, we coarse grain the initial dataset, effectively removing uninteresting local variation while maintaining the global structure of the dataset. Second, we use landmarking to quickly compute our diffusion potential coordinate system.

The initial coarse graining of the data is done to de-noise and downsample the input dataset while maintaining important global and local structures. By default, we downsample to 25,000 points using a fast hierarchical kmeans approach applied to randomized PCA dimensions. The centroids of these partitions are then passed along for diffusion potential calculation and condensation.

On this initial coarse-graining we compute the diffusion potential coordinates by employing landmarking as developed in [19]. Landmarking refers to the idea that instead of computing diffusion probabilities between every pair of points, we can compute diffusion probabilities from points to a well-chosen set of central “landmarks” that maintain the geometry of the data. From these landmarks, we can project back to the initial points, efficiently calculating diffusion potential coordinates for all points. The computation of the diffusion potential requires a powering of the diffusion probabilities, which has *O*(*n*^3^) complexity. With landmarking, this process speeds up by first using an *O*(*nk*) step for computing diffusion probabilities from the *n* points to the *k* landmarks, then only powering the smaller *k* × *k* diffusion matrix between the landmarks, reducing the overall complexity to *O*(2*nk*+*tk*^3^), where *t* is the number of diffusion steps. Finally, to speed up each condensation step, we use sparse multiplication to improve upon the *O*(*tk*^3^) complexity, speeding up each filtering step.

#### Selection of visualization scales and density aware PHATE rendering

While diffusion condensation reveals all levels of granularity in the data, we envision users to start with a coarse-grained level that offers a high-level summarization of the data before “zooming-in” to obtain finer detail on populations of interest. To offer this type of interactivity, we provide a method for selecting specific scales for visualization.

We reason that the salient levels of the representation for visualization must be stable levels which emerge during condensation, i.e., levels whose structure persists for several iterations. To find such levels we examine the gradient of points across successive condensation iterations and determine where the overall shift in data density from one iteration to the next is locally minimal (Figure 1B). Visualization of a granularity is achieved by the stress-minimizing optimization procedure provided by multidimensional scaling (MDS) on the filtered diffusion potential as done in [13]. Finally, to allow for more refined and detailed visualizations, we allow users to select data subsets to view at finer granularities identified by gradient analysis (Figure 1C).

**Figure 1:**
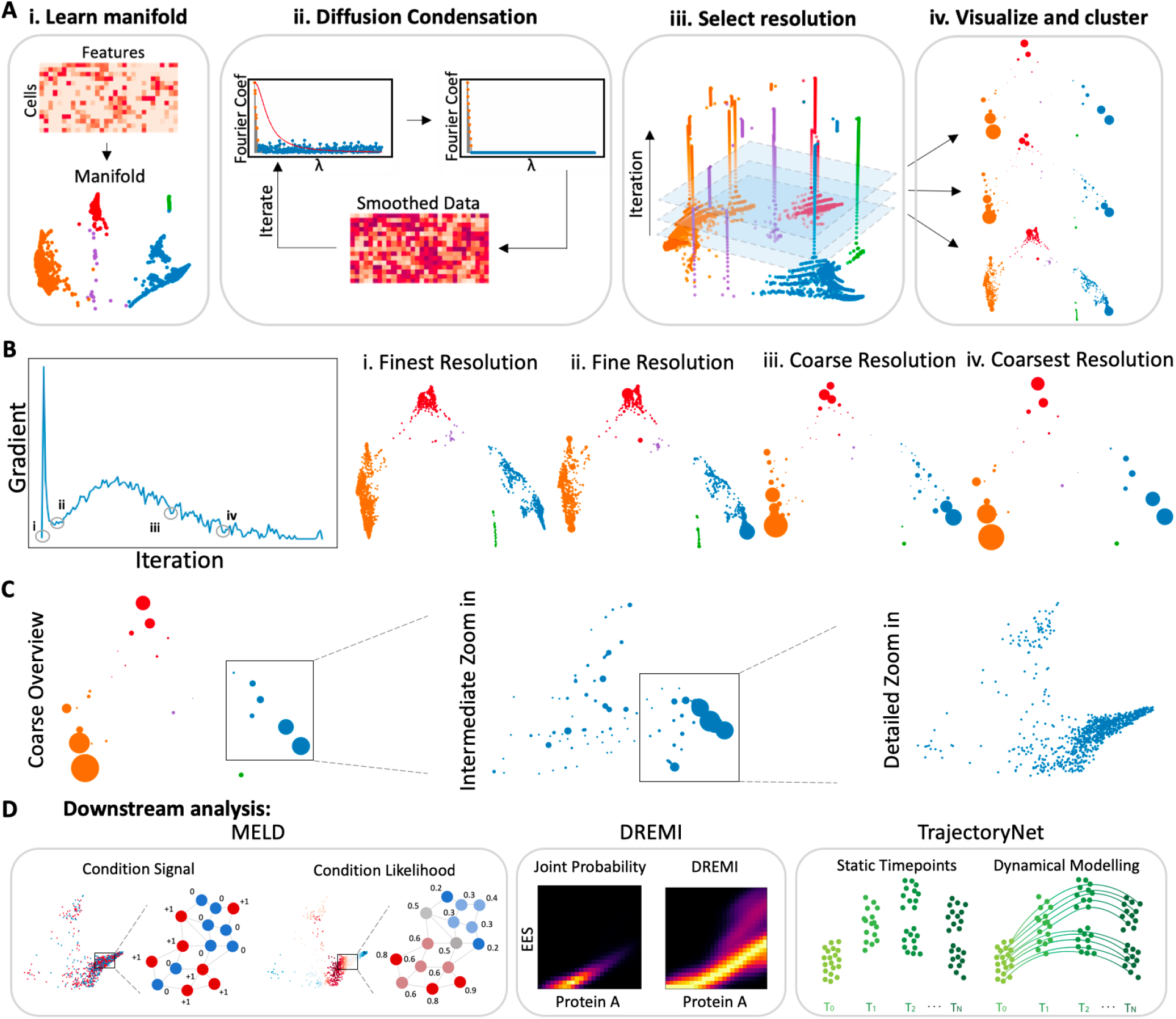
Overview of Multiscale PHATE algorithm. A. Multiscale PHATE process involves four successive steps. The first step (i) learns the manifold via diffusion potential calculation. The second step (ii) iteratively coarse grains the manifold construction through a fast diffusion condensation process. The third step (iii) involves the selection of salient granularities via gradient analysis before finally visualizing and clustering the manifold in the fourth step (iv). B. Gradient analysis identifies a range of scales for visualization. C. Multiscale PHATE allows for high level overviews of data as well as finer grain zoom ins of data subsets for additional detail. D. Multiscale PHATE abstractions of data are amenable to downstream analyses with algorithms like MELD [14], DREMI [15] and TrajectoryNet [16].

At each iteration, diffusion condensation merges points that are closer than a preset threshold distance. The PHATE rendering at each level is computed on the condensed diffusion potential where each point in the visualization is sized by the number of points that are merged to form that condensed point. This visualization strategy creates a naturally clustered and density-aware embedding.

#### Selection of clusters based on MELD mortality likelihood score to infer clinical associations

Multiscale PHATE abstracted data is amenable to many downstream computational analysis tools like MELD [14] for comparative analysis, DREMI [20] for computing mutual information between markers within subpopulations, and TrajectoryNet [21] for modeling dynamics of data (Figure 1D). In particular, to identify populations that are differentially enriched between experimental conditions or patients, we combine our powerful multigranular clustering approach with MELD [14] (Supplementary Figure 1B). MELD creates a joint graph of the samples being compared, and returns a relative likelihood that quantifies the probability that the each cell state in the graph is more likely in the control condition (which corresponds here to patients with positive outcome) or experimental condition (which corresponds here to patients with adverse outcomes). This likelihood score is found by first computing a cell-cell graph before creating an indicator signal for each of the two conditions. In our work, we often use patient clinical outcomes as the condition of origin. Then MELD smooths, or low-pass filters these signal over the cell-cell graph using the heat kernel to calculate the conditional density estimate of each condition over the cellular manifold. The density estimates of both conditions are then inverted via Bayes formula to create a relative likelihood of the condition given the cell. This likelihood score highlights regions of the manifold enriched in different conditions.

Finding a clustering method that matches the level of granularity of relative likelihood is a difficult problem, requiring the computationally complex vertex frequency clustering solution proposed previously [14]. However, Multiscale PHATE offers an alternative, less expensive solution as one of the granularities identified by diffusion condensation matches the clusters revealed by the MELD likelihood score. Combining the likelihood signal with our multigranular analysis, we can identify populations that are associated with particular outcomes with greater accuracy than other methods (Supplementary Figure 3B).

#### Construction of patient manifold through multiresolution cluster evaluation

After creating a cellular manifold by integrating hundreds of patients samples, it is critical to understand how similar or different each of these patients are from one another. Uncovering sample level density variations along the cellular manifold can be a powerful strategy to identify patient clinical states that are similar or dissimilar from one another. Previously, optimal transport approaches were used to construct a manifold of samples from single cell measurements systems [22]. Recently, these optimal transport based techniques have been implemented on hierarchical trees allowing for multiresolution comparison of sample density variations [23]. Furthermore, in [24] it was shown that multiscale smoothing of data distribution can be used to approximate the Earth-mover distances between them, which are closely related to optimal transport and essentially measures the energy required to transform one data distribution to another. Intuitively, this can be understood as computing how difficult it would be to change the underlying cellular state of one patient to that of another patient. Inspired by these techniques, we have developed a multiresolution evaluation system for determining sample level density variations to build a manifold of patients based on differences in their underlying cellular states. Until now, only two levels of the condensation tree have ever been evaluated simultaneously, typically one highlighting clusters that arise at a coarse level while another visualizing the data at a finer level. However, the groupings of points across multiple scales can provide rich information that can be harnessed by simultaneously evaluating cluster level information at multiple granularities.

With the goal of creating a manifold of patients, where each point represents a unique patient sample and distances between points represent how similar or different the underlying samples are in their cellular states as measured by flow cytometry, we evaluate clusters at multiple levels of the condensation tree. First, we identify all clusters at a particular level of the condensation tree. For a particular patient sample, we identify the proportion of the sample’s cells that fall within each of these clusters, creating a vector of cell proportions. This process is repeated for every sample, creating a set of features for every sample at a single resolution. This process is repeated for all samples across multiple resolutions to create an even richer, multiscale set of features related to prior work in multiresolution optimal transport [23]. This high dimensional multiscale feature matrix can then be embedded with PHATE for visualization.

### 4.2 Comparison of Multiscale PHATE to other visualization and clustering tools

#### Multiscale PHATE embeddings preserve local and global distances better than other multiscale visualization approaches

In order to quantify the quality of our dimensionality reduction strategy compared to other multiscale implementations of established visualization tools, we computed DeMAP scores [13] on embeddings of a variety of splatter simulated datasets [26]. While there is information loss in any sort of dimensionality reduction technique, an ideal embedding should capture as much local and global distance information as possible. To judge both local and global distance preservation, DeMAP quantifies the ability of an embedding to preserve ground truth manifold distances, also known as geodesic distances, in a low dimensional visual representation.

To compare the performance of Multiscale PHATE against six other dimensionality reduction tools at a particular granularity, we simulated scRNAseq data with Splatter [26]. Splatter uses a parametric model to simulate ground truth scRNAseq data with either cluster structure or path, also known as trajectory, structure. Splatter adds noise to this ground truth data, either by increasing dropout or by increasing biological variation, to simulate realistic noisy scRNAseq data on which different visualization and denoising tasks can be tested.

In our comparison, we fed increasingly noisy data to Multiscale PHATE and identified a salient level of granularity through gradient analysis for each run. After visualizing this level of granularity with Multiscale PHATE, we downsampled the noisy and ground truth datasets to this particular scale in order to compute embeddings with other dimensionality reduction algorithms at the same resolution. We computed DeMAP scores on each of these embeddings, comparing the geometrically downsampled ground truth splatter data to the embedding of the same granularity. We repeated this simulation on a diverse set 10 datasets with clusters and 10 datasets with paths. As we increased the degree of dropout and biological variation, Multiscale PHATE captured ground truth structure of the data more robustly than any other method, outperforming all other methods in 18 of the 20 comparisons (Figure 2C). From these experiments, we conclude that Multiscale PHATE visualizes coarse grained data better than other visualization techniques.

**Figure 2:**
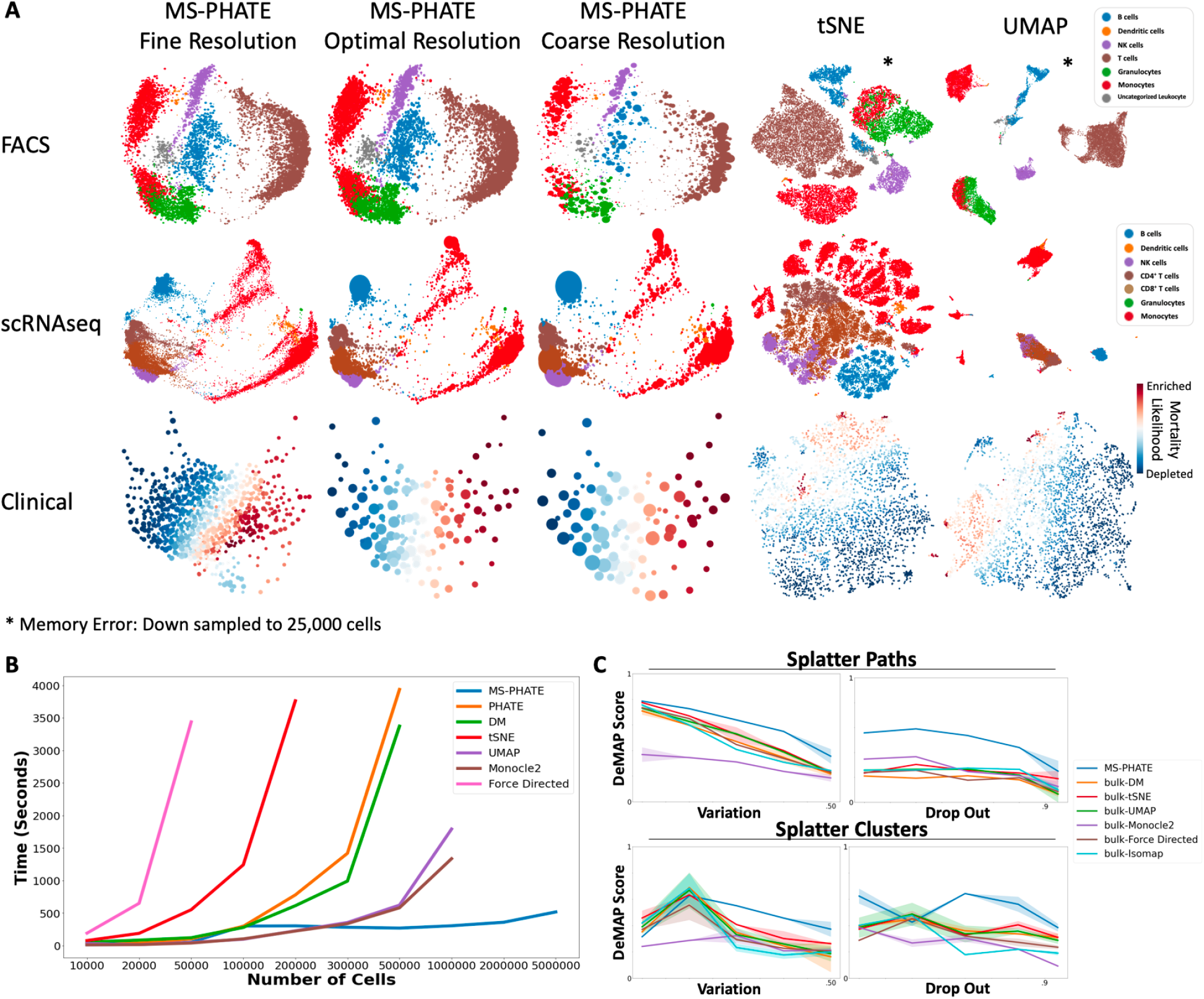
Comparison of Multiscale PHATE with other dimensionality reduction tools. A. Visual comparison of Multiscale PHATE with other dimensionality reduction tools across a range of data types, including 22 million PBMCs measured by flow cytometry [25], 49,942 PBMCs by scRNAseq [10] and 2,135 patients admitted to YNHH by demographic and lab clinical variables. B. Comparing run time across visualization techniques on increasingly high dimensional flow cytometry data. C. Quantitative comparison between embeddings produced by Multiscale PHATE and other hierarchical visualization strategies. Comparisons were evaluated using DeMAP with increasing levels of 2 different types of biological noise, drop out and variation, as well as on data with different structures, clusters and paths.

In addition to quantitative evaluation of Multiscale PHATE, we can also assess its performance qualitatively. When visualizing high dimensional cellular data, we would expect a good visualization technique to embed all cells of a single type close to one another and cells of different types far apart. When comparing our method to other dimensionality reduction techniques, it becomes clear that Multiscale PHATE more accurately maintains distances between similar and dissimilar cell types (Figure 2A). When comparing monocyte clusters derived from flow cytometry data across a range of visualization techniques, we see that across all Multiscale PHATE granularities, monocytes are embedded close to one another but form distinct and distant clusters in tSNE and UMAP visualizations. When evaluating embeddings of single cell transcriptomics data [10], these monocytes are preserved in three groups by UMAP and across a range of Multiscale PHATE granularities, however are broken into over a dozen groupings by tSNE. Similarly, when embedding patient clinical data, Multiscale PHATE visualizes patients with similar clinical outcomes close to one another. These relationships are broken into more distant clusters by tSNE and UMAP embeddings.

Finally, it is clear that Multiscale PHATE is far more scalable and reproducible than other visualization techniques. While Multiscale PHATE is able to embed 22 million cells measured by flow cytometry in less than 20 minutes, users are forced to downsample this dataset to just 25,000 cells in order to create UMAP or tSNE visualizations (Figure 2A). When comparing run times between different techniques, it becomes clear that Multiscale PHATE is able to rapidly scale to millions of cells, successfully embedding 5 million cells in less than 10 minutes, while the next most scalable technique, Monocle2, can only embed 500,000 cells in a comparable time (Figure 2B). Finally, Multiscale PHATE is highly reproducible. A common issue with UMAP and tSNE, which shift clusters randomly from run to run, is solved by Multiscale PHATE, which can faithfully create the same embedding across multiple runs (Supplementary Figure 2B).

#### Multiscale clusters more accurately capture established groupings of data in synthetic and real biological datasets

While data scientists sometimes think of neighborhood embeddings such as tSNE or UMAP as clustering methods, since they visualize data in a manner reminiscent of clusters, such methods do not explicitly return groupings of the data. Instead, the associated data has to be processed by a clustering algorithm, such as *K*-means [27] or Louvain [28], to produce groups. In contrast, Multiscale PHATE is truly both a visualization and clustering algorithm, as the condensation process effectively returns groupings of the data at all levels of granularity.

In order to quantify the clustering accuracy of Multiscale PHATE on increasingly noisy and multigranular data, we simulated two and three-layer hierarchical stochastic block models (SBM) (Figure 3A, Supplementary Figure 3A). In these models, a graph is constructed in which there are coarse grain clusters, each of which could be further broken down into increasingly granular clusters. In order to compare all clustering techniques across a range of noise levels, increasing amounts of random gaussian noise is added to the edge weights of the graph. At each level of noise, cluster labels are computed with multiple clustering tools: Multiscale PHATE, Louvain [28], Leiden [29] and single linkage hierarchical clustering [30]. For each clustering tool, the entire tree of cluster assignments is evaluated against coarse and fine grain ground truth cluster labels using Adjusted Rand Index (ARI), with the top score preserved. These comparisons are run on a range of noise levels and replicated 10 times across a range of initial hierarchical SBM edge weights in both the two-layer and the three-layer models. Across both models, Multiscale PHATE performed superior to other hierarchical clustering techniques in 35 of the 42 comparison conditions (Figure 3A, Supplementary Figure 3A), with only poorer performance at the finest granularity of the 3 layer SBM. From these experiments, we conclude the Multiscale PHATE is superior at clustering noisy simulated data than other leading multigranular clustering tools.

**Figure 3:**
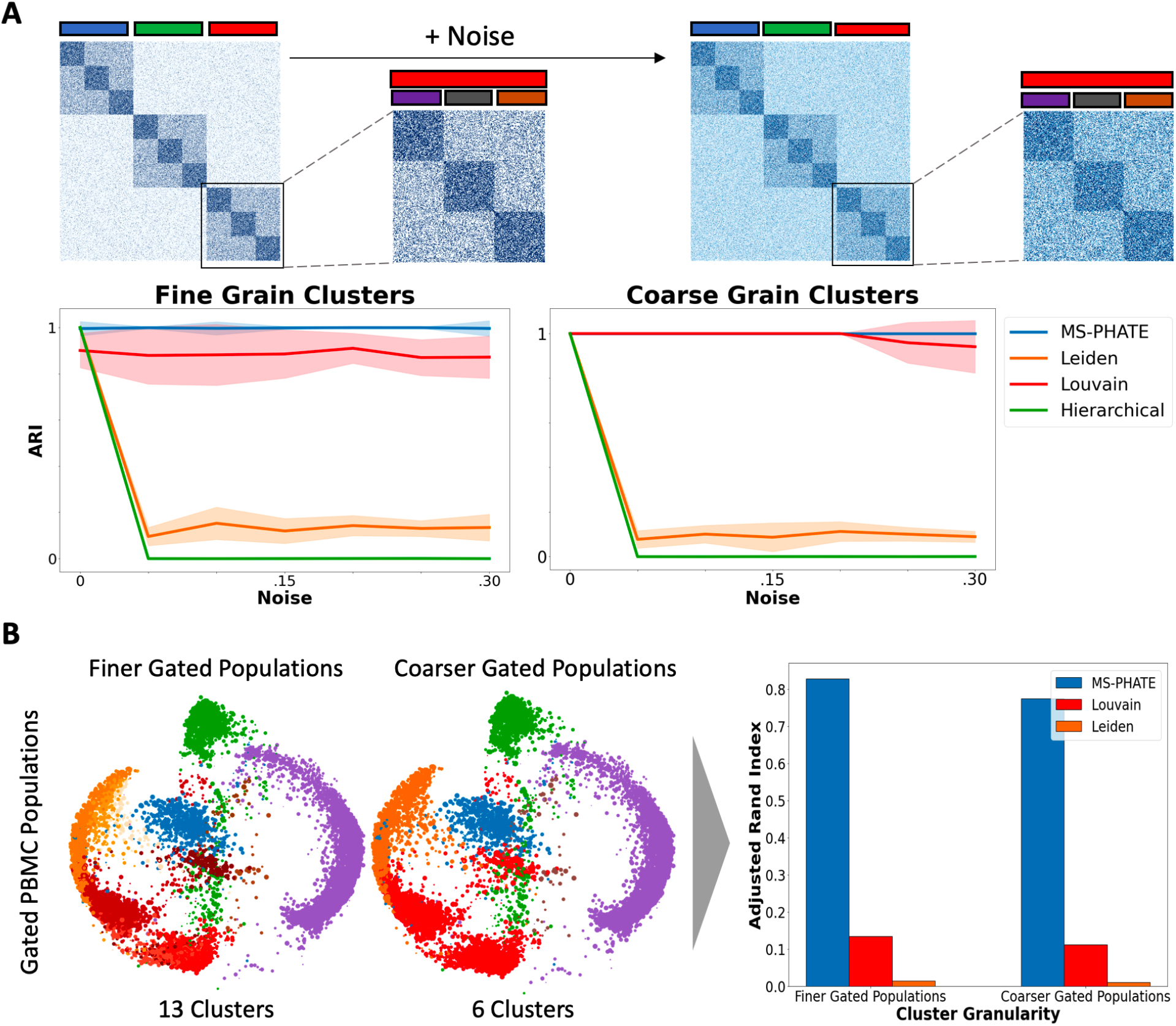
Comparison of Multiscale PHATE with other Clustering techniques. A. Schematic of the hierarchical stochastic block model we generated for multigranular cluster comparisons. For each method, increasing amounts of random Gaussian values were added to the adjacency matrix of stochastic block model to simulate increasing amounts of noise. As the model was constructed with known clusters at multiple scales, we computed Adjusted Rand Index (ARI) between each algorithm’s predicted clusters and the known clusters across coarse and fine granularities. B. Comparison of multiple clustering approaches on flow cytometry data where cell types and subtypes have been identified through gating analysis. Clusters identified by different approaches were compared to gated populations using ARI.

Traditional methods to analyze flow cytometry data involve individually establishing thresholds across the fluorescent signals from each targeted marker on a given cell based on the controls. By iterating similar “gates” across a number of markers, an investigator may identify increasingly specific cells within a heterogeneous population. Individual gates for each marker may be validated by comparing against rigorous controls including isotype controls, fluorescence minus one controls, and biological controls. These methods can distinguish true fluorescent signal from non-specific signal or biological noise, respectively. This robust approach remains the gold standard for the analysis of flow cytometry data [31]. In order to show that Multiscale PHATE is able to identify known cell types, we computed multiscale clusters on a dataset with cell types established by traditional gating analysis. Taking the cell population labels as identified conventionally, we computed ARI scores on clustering outputs from a range of clustering techniques at multiple granularities. Across both fine and coarse grain clusters, Multiscale PHATE computed clusters that more faithfully represented the underlying known biological cell types (Figure 3B).

### 4.3 Multiscale PHATE analysis of 251 SARS-CoV-2 patient blood samples reveals subsets of cells associated with mortality

One hundred sixty eight patients with moderate to severe COVID-19 [32] were admitted to YNHH and recruited to the Yale IMPACT (Implementing Medical and Public Health Action Against Coronavirus CT) study. From each patient, blood samples were collected across multiple timepoints to characterize patient cellular responses across the spectrum of disease. In total, the composition of peripheral blood mononuclear cell (PBMC) was measured by flow cytometry on 251 samples. Finally, clinical data was extracted from the electronic health record corresponding to each biosample timepoint to allow for clinical correlation of findings (see Methods). In this analysis, we define poor or adverse outcomes as patients who died from infection, while good outcomes as patients who survived. In order to analyze over 54 million cells characterized across 4 different sets of flow marker panels, we applied Multiscale PHATE to identify subsets of peripheral blood mononuclear cells (PBMCs) associated with mortality and survival.

#### Key dysfunctional myeloid, granulocyte and B cell subsets are enriched in patient who die from infection

To explore the role of individual PBMC cell types in disease pathogenesis, we examined 22 million cells measured on a myeloid-centric flow cytometry panel from 210 patient samples suffering from COVID-19. Using cell type specific marker staining, we characterized Multiscale PHATE clusters (Figure 4A). Using MELD and the mortality outcome for each patient in our cellular state space, we were able to compute the mortality likelihood score, which identified cellular states enriched in patients who die from infection (darker red) or patients that survive (darker blue) (Figure 4B). When mapping these scores onto cluster labels, we found that the three populations most enriched in mortality were granulocytes (CD16^+^SSC^*hi*^), B cells (CD19^+^), and monocytes (CD14^+^) while the population most enriched in survival was T cells (CD3^+^) (Figure 4C).

**Figure 4:**
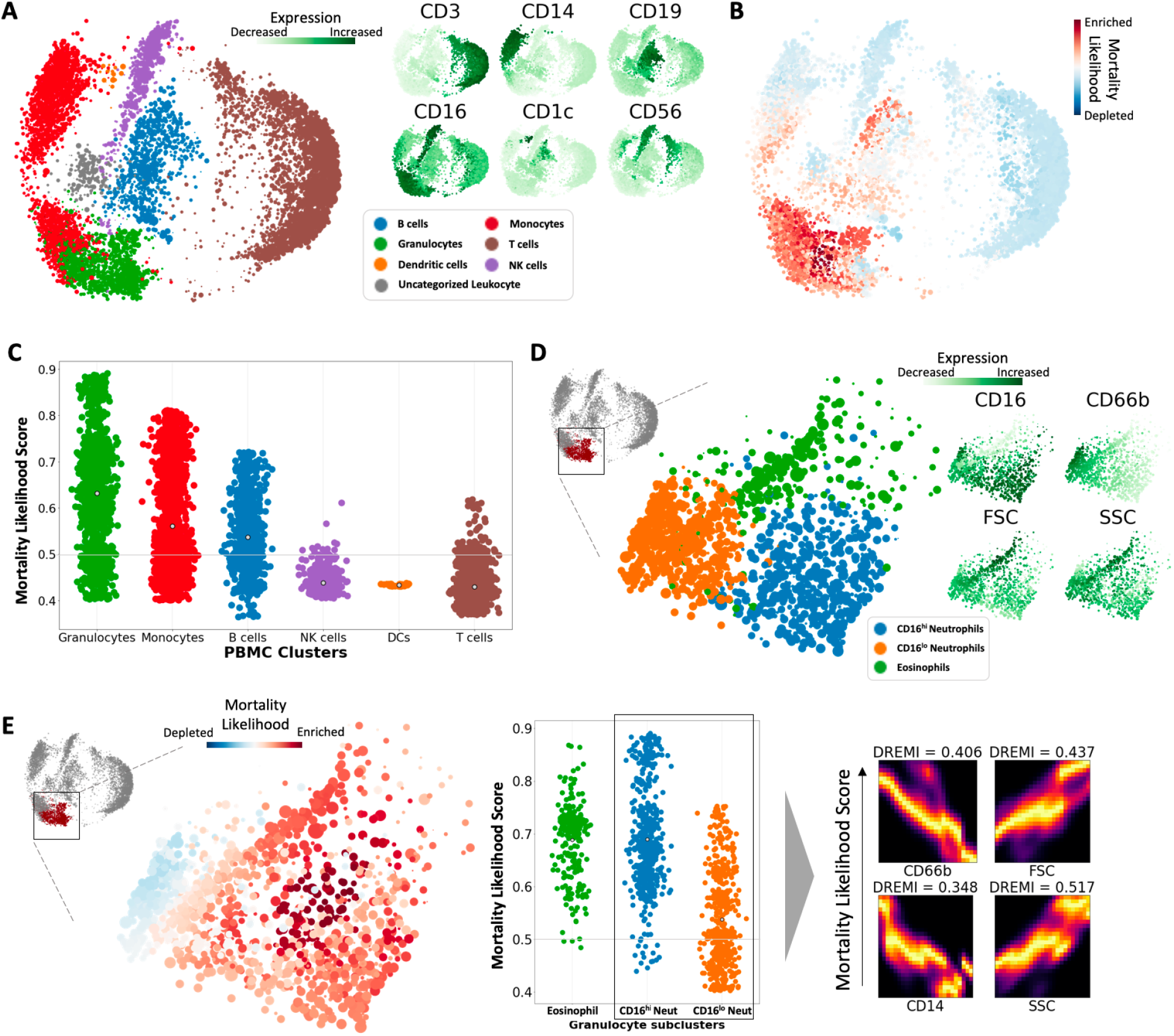
CD16^hi^ CD66b^lo^ Neutrophil subset enriched in patients who die from COVID-19. A. Multiscale PHATE visualization of PBMCs identifies all major cell types based on cell type specific markers. B. Visualization of mortality likelihood score computed by MELD. C. Visualization of mortality likelihood score organized by cell type reveals enrichment of granulocytes, monocytes and B cells in patients who die from COVID-19. D. Zoom in of granulocyte population identifies subsets of neutrophils and eosinophils based on expression of known markers. E. Visualization of mortality likelihood score in granulocyte population identifies CD16^hi^ neutrophils enriched in patients with worse outcomes. Key associations between markers and mortality likelihood scores in neutrophils computed by DREMI and visualized with DREVI.

#### Resting population of circulating neutrophils enriched in with patients who die from COVID-19

We zoomed in on the granulocyte population and identified CD16^*hi*^ neutrophils, CD16^*lo*^ neutrophils and eosinophils based on the expression of CD16, CD66b, granularity by side scatter (SSC) and size by forward scatter (FSC) (Figure 4D). After mapping our mortality scores onto this granulocyte population we found that the CD16^*hi*^ neutrophils were enriched in patients who died from infection. In order to identify which cellular markers beyond CD16 were most correlated with mortality in neutrophils, we computed DREMI between protein expression and mortality likelihood scores in both neutrophil subsets. We identified that while CD14 and CD66b were negatively correlated with mortality, increased FSC and SSC were both strongly positively correlated with mortality in neutrophils, indicating that CD16^*hi*^CD66b^*lo*^ neutrophils were enriched in patients that died from COVID-19. Based on the PBMC isolation protocol used (see Methods) neutrophils obtained were by definition Low-Density Neutrophils, containing both the mature and immature subsets. Considering the sensitivity of CD16 expression, high CD16 in our cohort was most likely indicative of a mature population that has not responded to an activating stimulus [33–35]. Neutrophils from patients with worse disease also expressed less CD66b; in contrast, an increase in surface expression of CD66b occurs following degranulation [36]. The combination of high complexity, high CD16 expression, and low CD66b expression suggests a resting population of circulating neutrophils present in patients with lethal disease.

#### CD14^−^CD16^*hi*^HLA-DR^*lo*^ monocyte subset associated with patient mortality

In order to identify monocyte subsets implicated in disease, we zoomed in to the monocyte population and identified major subtypes based on the expression of markers CD16 and CD14 (Supplementary Figure 4A). The combination of these markers allowed us to distinguish between CD14^+^CD16^−^ monocytes, CD14^+^CD16^*int*^ monocytes and CD14^−^CD16^*hi*^ monocytes. After mapping our computed mortality likelihood scores onto this population, we identified that CD14^−^CD16^*hi*^ monocytes were the most strongly enriched in severe infection, followed by CD14^+^CD16^*int*^ monocytes (Supplementary Figure 4B). These findings agreed with published observations as others have also noted an influx of CD14^+^CD16^*int*^ and CD14^−^CD16^*hi*^ monocytes in the lungs of patients with severe disease [11, 37, 38]. Furthermore, across all monocytes, CD16 was positively correlated with mortality while CD14 and HLA-DR were correlated with survival, identifying a distinct CD14^−^CD16^*hi*^HLA-DR^*lo*^ population of monocytes enriched in mortality. The loss of HLA-DR has been previously shown in monocytes from COVID-19 patients [39]. Monocytes expressing HLA-DR can serve as antigen-presenting cells to shape the adaptive T cell response, but monocytes in this cohort, expressing reduced amounts of HLA-DR, would likely have very limited capacity to prime effector T cell responses. Interestingly, a similar phenomenon occurs in sepsis patients, as well, and is indicative of worse outcomes [40–42]. In this setting, elevated levels of IL-10 have been linked to decreased HLA-DR on monocytes [43, 44]. As COVID-19 patients also present with significantly elevated levels of IL-10 in circulation, a similar mechanism may be at play here [39, 45].

#### Multiscale PHATE identified plasmablast population associated with mortality

There has been a persistent interest in the role of B cells during disease due to their potential to generate neutralizing antibodies. In our broad PBMC analysis, however, B cells were among the most enriched populations in severe outcomes (Figure 4C). In order to explore B cells in greater detail, we processed 154 patient samples on a B cell specific flow cytometry marker panel. Analyzing these cells by Multiscale PHATE granted us an unbiased, granular look at B cells subsets which would otherwise be difficult by traditional two-dimensional gating, popular for flow cytometry analysis. These subsets include transitional B cells (IgD^+^IgM^+^CD27^−^/CD38^+^CD24^+^), naïve B cells (IgD^+^IgM^+^CD27^−^/CD38^−^), switched (IgD^−^IgM^−^CD27^+^) and unswitched memory B cells (IgD^+^IgM^+^CD27^+^), activated B cells (IgD^−^CD138^−^CD86^+^HLADR^+^), and antibody secreting cells (CD138^+^CD38^+^) (Supplementary Figure 4C). After identifying these major cell types, we computed mortality likelihood scores to identify B cell subtypes implicated in mortality. Interestingly, the most enriched cell type in patients with adverse outcomes was a subset of the antibody secreting population defined by CD86^*lo*^HLADR^−^/CXCR3^+^, also known as plasmablasts. Meanwhile the cell types most enriched in patients with good outcomes was a subset of late activated mature B cells defined by CD86^+^ (Supplementary Figure 4D). Despite the well-described protective roles of circulating antibodies, these results are consistent with earlier findings from COVID-19 patients, which discuss the potential role of B cells in disease pathogenesis [46–48]. Given the abundance of circulating IL-6 in COVID-19 patients, it is possible that in this setting, IL-6 skews B cell differentiation into antibody secreting cells [49]. Considering the lack of mutations in neutralizing, anti-SARS-CoV-2 antibodies, skewing toward antibody secreting cells may come at the expense of transit through the germinal center, thus producing a less potent or non-specific antibody response [50].

#### Key pathogenic T cell subsets are enriched in patients who die from infection

Although T cells collectively were enriched in patients who recovered from infection (Figure 4C), there are a diverse set of T cell subsets which have been implicated in severe disease pathogenesis. In order to identify functional T cell subsets enriched in patients who died from COVID-19, we applied Multiscale PHATE to 22 million T cells measured on a cytokine-specific flow cytometry panel. After identifying salient levels of granularity for downstream analysis, we identified both CD4^+^ and CD8^+^ T cell subsets at coarse granularity (Figure 5A).

**Figure 5:**
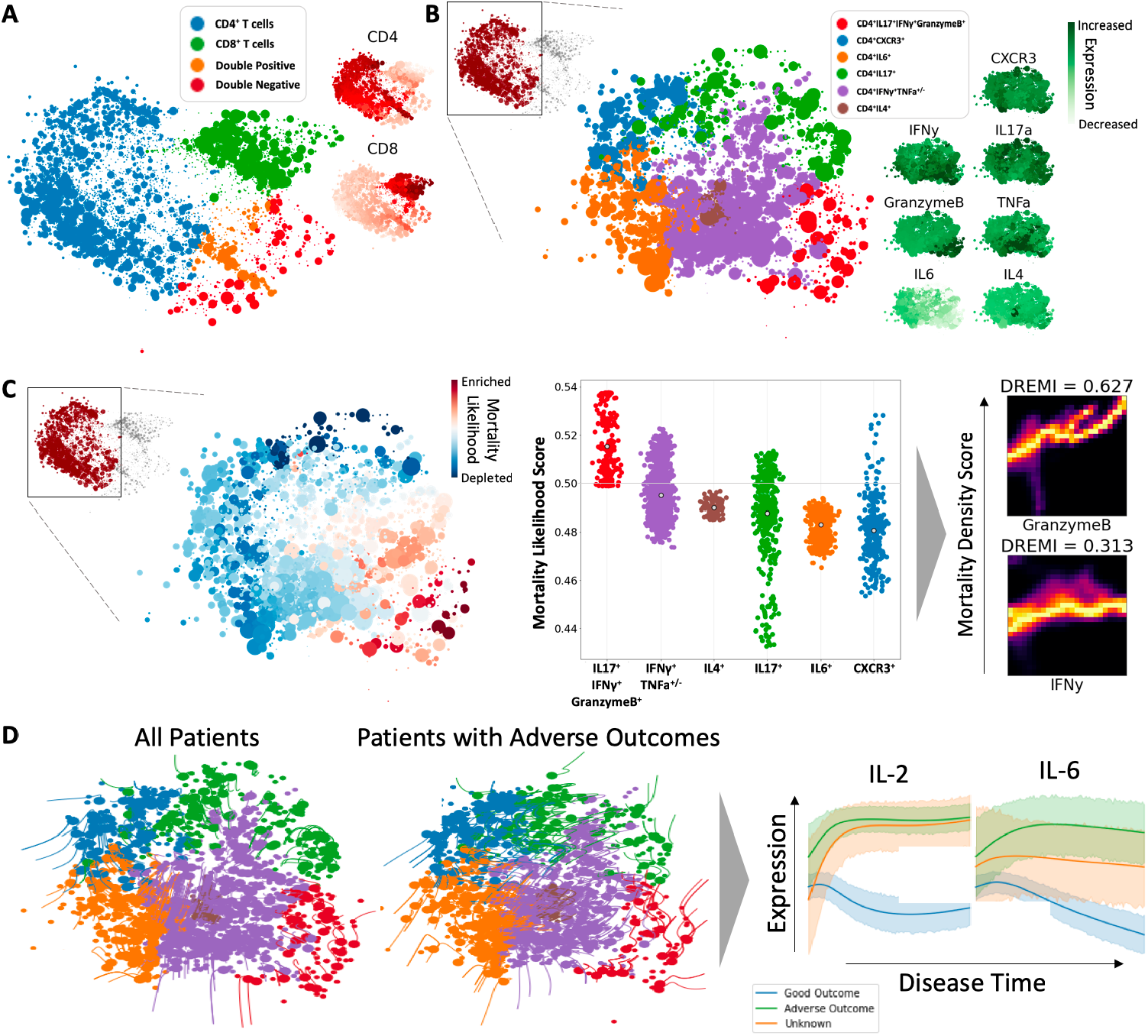
Multiscale PHATE identifies Th17 subset enriched in patients who die from COVID-19. A. Multiscale PHATE visualization of T cell focused cytokine panel identifies broad T cell subtypes. B. Zoom in of CD4^+^ T helper cells identifies subsets based on expression of functional markers. C. Visualization of mortality likelihood score identifies IFNγ^+^ GranzymeB^+^ Th17 cell enrichment in patients with poor outcomes. Key associations between markers and mortality likelihood scores are computed by DREMI and visualized with DREVI. D. TrajectoyNet run on CD4ff T cells from all patients and patients who died from infection identifies condition-specific cellular dynamics. Tracking the expression of key markers overtime, TrajectoryNet identifies mortality dependent differentially expressed markers in CD4^+^ T cells overtime.

#### Fine grain analysis of protective T cell population helps identify pathogenic IFNγ^+^ GranzymeB^+^ Th17 subpopulation

Using Multiscale PHATE’s zoom and cluster capabilities, we were able to visualize the CD4^+^ T cells and subdivide the cells into functional subsets using functional markers, IFNγ, IL-17, and IL-4 (Figure 5B). Interestingly, in our dataset, we identified two different subsets of CD4^+^ IL-17 producing T cells classically known as Th17 cells, one co-producing GranzymeB and IFNγ and one staining negative for both markers. Finally, we classify two final subsets based on the expression of IL6 and CXCR3 (Figure 5B). To identify cell types enriched in mortality, we computed a mortality likelihood score. By organizing our scores by T helper subset, it became clear that the Th17 subset co-producing IFNγ^+^GranzymeB^+^ were enriched in patients who died from infection. Furthermore, GranzymeB and IFNγ were positively associated with mortality likelihood on DREMI analysis across all CD4^+^ T cell subtsets (Figure 5C).

While Th17 cells can play protective roles [51], IFNγ^+^GranzymeB^+^ Th17 cells have been associated with tissue damage. In a model of murine autoimmune encephalomyelitis, a discrete subset of IFNγ^+^GranzymeB^+^ Th17 cells caused significantly worse disease than traditionally activated Th17 cells [52]. Previous literature had also observed that high levels of circulating IL-17 produced from IFNγ^+^GranzymeB^+^ Th17 cells could drive a strong pro-inflammatory immune response and promote neutrophil expansion. Likewise, recent reports have indicated the harmful contribution of neutrophils and neutrophil extracellular traps (NETs) in SARS-CoV-2 infections [53–55]. This influx of neutrophils can be further exacerbated by the virus-induced loss of ACE2 [56], and cumulatively, these events have the potential to trigger ARDS, as seen in COVID-19 patents. However, what regulates neutrophil recruitment, survival, and subsequent NET release during the disease has not been definitively identified in COVID-19. Interestingly, patients with adverse outcomes in this cohort demonstrated an enrichment in IFNγ^+^GranzymeB^+^ Th17 cells, as well as CD16^+^ neutrophils. We posit that IFNγ^+^GranzymeB^+^ Th17 cells in our cohort may precipitate these pathogenic effects via IL-17 secretion. While our flow cytometry data was limited to identifying cells in circulation, sequencing data from upper respiratory tracts of COVID-19 patients observed Th17 cells in the airways, as well [57, 58]. Either in the lungs or in circulation, IFNγ^+^GranzymeB^+^ Th17 may influence neutrophil activity by inducing IL-8 release from airway epithelial cells or G-CSF from microvascular pericytes [59–61]. It was recently shown that MAIT cells comprise a substantial portion of IL-17 expressing cells in the upper respiratory tracts of COVID-19 patients; consequently, secretion of IL-17 in the lung may not be primarily confined to the Th17 compartment. The two may also act synergistically as MAIT cells have been shown to promote the recruitment of activated CD4^+^ T cells to the lungs during pulmonary infection [62].

To understand cellular dynamics from our multiscale distilation, we ran TrajectoryNet [21] on our CD4^+^ T cell coarse grained embedding as well as on the underlying markers. With this analysis we identified diverging trajectories when comparing all patients with patients who experienced adverse outcomes (Figure 5D). When comparing trajectories from these patient populations, we can identify a clear set of Th17 cells from patients with adverse outcomes shifting towards the IFNγ^+^GranzymeB^+^ co-producers. This indicates, that in patients with adverse outcomes, potentially homeostatic Th17 cells acquire a pathogenic phenotype. By integrating TrajectoryNet with Multiscale PHATE, we also show that we can identify biomarkers that are predictive of disease outcome. We identify that IL-2 and IL-6 expression by CD4^+^ T cells is differentially based on patient outcome over time, with patients who die expressing higher levels of IL-2 and IL-6 over time and patients who survive expressioning lower amounts of both (Figure 5D).

#### Hyperactivated CD8^+^TIM3^+^HLA-DR^+^PD1^+^ TEMRA cells, expressing GranzymeB enriched in patients who die from COVID-19

In acute viral infections, CD8^+^ T lymphocytes play a critical role in the clearance of virus [63]. By the directed secretion of Granzyme B, these effectors may rapidly kill virally-infected targets [64]. In order to characterize the role of CD8^+^ T cell subsets in disease, we zoomed in on CD8^+^ T cells in our cytokine-focused T cell panel. Using the expression of cell surface markers and cytokines, we identified three major subsets, one producing GranzymeB, one producing IFNγ and one producing TNFα (Supplementary Figure 5A). After mapping mortality likelihood scores onto the CD8^+^ subpopulation, it became clear that the GranzymeB^+^ population is most enriched in mortality with GranzymeB expression being highly associated with mortality in CD8^+^ T cells (Supplementary Figure 5B). These findings are consistent with a previous study of SARS-CoV-2 infected patients that observed an association between the enrichment of CD8^+^ T cells expressing high amounts of GranzymeB and increased disease severity [65]. Despite the protective role of GranzymeB in other viral infections, our data and others indicate that its excess may lead to worse outcomes, including mortality. This possibility is supported by early findings of GranzymeB^+^ CD8^+^-induced tissue damage in different murine models of respiratory viral infections [66, 67] or from clinical studies [68]. In these early studies from mice, pathogenic GranzymeB^+^ CD8^+^ T cell activity manifested in the presence of extremely high viral loads or in the absence of other lymphocytes and antigen-specific antibodies. Likewise, all of these factors are present in COVID-19 patients-high viral loads, lymphocytopenia, and ineffective antibody responses - which permits the emergence of this hyper-activated CD8^+^ population associated with more severe disease. To gain additional insight on which discrete subset of CD8^+^ T cells may be the source of GranzymeB, we performed detailed surface staining of all T cells.

We analyzed 208 patient samples using a flow cytometry panel containing markers indicative of T cell subset identity and activation status. After identifying the ideal granularity to analyze the data, we identified CD4^+^, CD8^+^ and double positive T cell subsets (Supplementary Figure 5C). Zooming in to the CD8^+^ subset, we identified a range of activation states based on the expression of key markers: Effector (TIM3^+^PD1^+^/CD45RA^+^), Follicular (CD45RA^−^/CCR7^−^CD127^−^/CXCR5^+^PD1^+^), Memory (CD45RA^−^/CCR7^+^ or CCR7 ^−^ CD127^+^), Naive (CD45RA^+^/CCR7^+^CD127^+^) and T effector memory cells re-expressing CD45RA (TEMRA) (CD45RA^+^/CCR7^−^CD127^*lo*^) (Supplementary Figure 5D). After computing MELD mortality likelihood score, we identified that the TEMRA population displayed the most enrichment in severe infection. Furthermore, across all CD8^+^+ T cells, activation state markers PD1, TIM3, HLA-DR and CD45RA were also positively correlated with mortality on DREMI analysis, while markers like CD127 and CCR7 were negatively correlated with mortality (Supplementary Figure 5E). Our findings here are in agreement with contemporaneous studies of SARS-CoV-2 patients [46, 65, 69]. Cumulatively, our data correlate mortality with a hyper-activated CD8^+^ T cell response in the form of CD8^+^CD45RA^+^TIM3^+^HLA-DR^+^PD1^+^ TEMRA cells, likely expressing GranzymeB.

### 4.4 Multiscale patient manifold construction reveals potential mechanisms of disease

Here, we show that Multiscale PHATE-derived clusters across multiple scales form a rich set of feature descriptors for patients measured in single cell modalities. Although, the purpose of measuring single cell data is indeed to derive features in the form of cells, patients can be hard to compare and analyze at this level. Common approaches simply compare cluster proportion or averaged expression levels across patients. However, since Multiscale PHATE creates cellular groupings at multiple granularities, we can derive a rich summarization of patients across scales.

We create a patient-embedding using cluster proportions from several levels of the condensation tree of the myeloid-focused flow cytometry using our patient manifold approach (Figure 6A). Briefly, the proportion of a patient’s cells that belong to clusters at several levels of the tree are used as a feature vector (see Methods). These patient descriptors are then embedded and visualized with PHATE [13]. The resultant embedding demonstrates that the patients lie on a low dimensional continuum or manifold themselves. When the patient embedding is colored by the manifold-based likelihood estimate of mortality outcomes, we see that the dominant progression in the data is indeed clinical outcome, with patients on the left enriched for good outcomes (darker blue) and patients on the right enriched for adverse outcomes (darker red).

**Figure 6:**
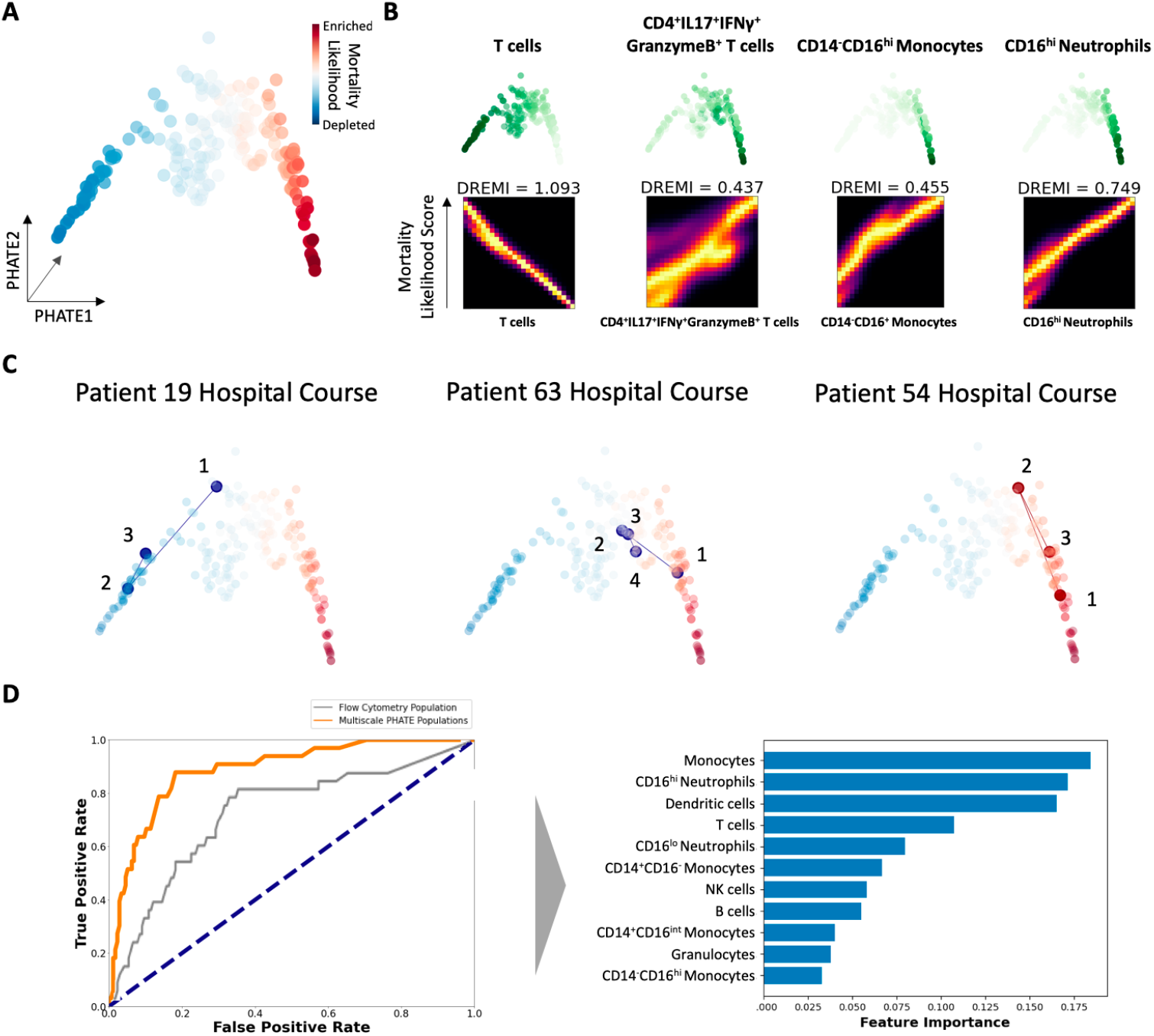
Patient manifold corroborates cellular states associated with of disease pathogenesis. A. Visualization of patient manifold via PHATE and mortality likelihood score based on patient outcomes computed via MELD. B. Visualization of key cell population enrichment trends over the manifold with associations computed by DREMI and visualized with DREVI. C. Tracing three patients’ hospital courses over patient manifold. Patients 19 and 63 were discharged while patient 54 died. D. Comparing predictability of patient mortality using random forest classifier on Multiscale PHATE identified populations and flow cytometry identified populations. Most predictive Multiscale PHATE clusters are ranked through feature importance analysis.

In order to associate previously identified cellular populations with outcome, we computed DREMI between these population proportions and mortality likelihood score. We identified that while T cells overall were negatively correlated with mortality, CD4^+^IFNγ^+^GranzymeB^+^ Th17 cell, plasmablasts, CD16^*hi*^ neutrophils and CD14^−^CD16^*hi*^ monocytes were all strongly positively associated with mortality (Figure 6B). These findings indicate that precipitous decline in T cells correlates with mortality, while subsets of neutrophils, monocytes and Th17 cells, all previously highlighted in our analyses, are increased in patients with adverse outcomes. Finally, we trace clinical states of three patients, 19, 63 and 54, across the patient manifold to determine if our construct accurately recapitulates patient trajectories. Surviving patients 19 and 63 had their clinical trajectories consistently go from the high mortality region to the low mortality region. In contrast, patient 54, who succumbed to disease, had a tortuous set of clinical states all of which mapped within the high mortality region (Figure 6C).

To identify if age, sex and other clinical variables were preferentially associated with mortality on our construction, we mapped these clinical variables onto the patient manifold. We found that patients who were more likely to experience poor outcomes were also more likely to be older, male, receive ventilatory support and have higher markers of inflammation (Supplementary Figure 6A). We ran DREMI analysis to find associations between these clinical variables and key cell types implicated in infection pathogenesis. We found that females and young individuals were more likely to mount a robust T cell response. This finding builds upon a body of literature that shows immune responses may differ across sex and age [70, 71]. Specifically, for SARS-CoV-2, this finding corroborates a separate analysis that found that activated T cells play a protective role in women but not as much in men [72]. Additionally, the negative relationship between age and T cell numbers has been extensively studied [73, 74]. Our analysis finds the same trend in our cohort of patients, who are in general older, predisposing them to requiring hospitalization. Epidemiological data analyzing large numbers of COVID-19 patients enumerate the significant contributions of age and sex for disease severity [75, 76].

In order to see if Multiscale PHATE-derived sub-populations could predict disease outcome, we combined the features of patients we identified in our myeloid-focused flow cytometry panel with clinical outcome to train a random forest classifier (see Methods). Using these abstracted features, we accurately predict outcome in 83.5% of cases. When performing a similar prediction task using flow cytometry gated populations, we were only able to predict outcome with 73.8% accuracy. Furthermore, we identified that monocytes, CD16^*hi*^ neutrophils and T cells were three of the top four cell types most predictive of eventual disease outcome in our classifier model (Figure 6D).

### 4.5 Multiscale clinical manifold construction highlights potential mechanisms of disease convalescence

Thus far, we have primarily used Multiscale PHATE to identify multiresolution structure in single cell flow cytometry data. We now showcase the utility of Multiscale PHATE on a different type of data: laboratory, clinical, and demographic data generated from routine clinical care of COVID-19 patients admitted to YNHH. With 18 clinical variables collected on 2,135 patients admitted to YNHH diagnosed with moderate to severe COVID-19, we created a multiscale embedding capturing patient states across mild, moderate and severe spectrums of disease. Patient outcomes at discharge were categorized by severity: discharge to home, discharge to rehabilitation for extended recovery, and discharge to hospice or death while in hospital. Using each of these outcomes, we computed likelihood scores with MELD corresponding to each outcome: survival likelihood score, extended recovery likelihood score and mortality likelihood score (Figure 7A).

**Figure 7:**
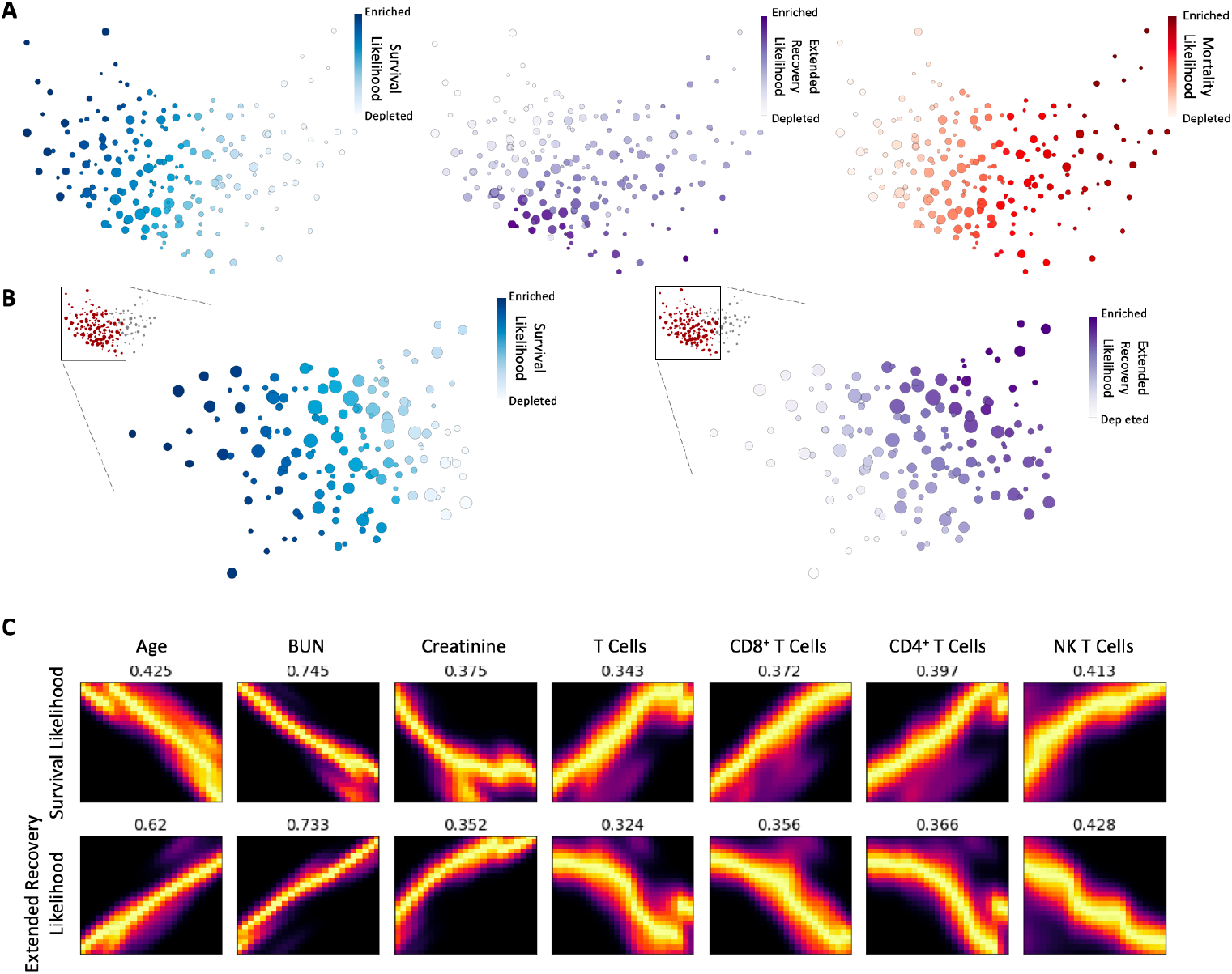
Multiscale manifold of patient clinical features identifies cell types associated with extended COVID-19 recovery phase. A. Visualization of Multiscale PHATE clinical manifold constructed on patient clinical features. Embedding is colored by likelihood scores based on patient outcomes computed via MELD. B. Zoom in on transition point between high extended recovery likelihood score and high survival likelihood score. C. Patient clinical features and flow cytometry identified cell populations associated with patient outcomes using DREMI and visualized with DREVI.

In order to understand how clinical features could inform outcomes, we computed DREMI and DREVI analysis between clinical features and each of our likelihood scores (Supplementary Figure 7A-B). As anticipated, markers of physiologic instability, such as decreased systolic blood pressure and increased respiratory rate, as well as systemic inflammatory markers, increased ferritin, procalcitonin and CRP, were associated with higher mortality. Beyond these general markers, however, several markers of organ dysfunction were also strongly associated with mortality. Specifically, kidney dysfunction, as measured by blood urea nitrogen (BUN) and creatinine, as well as liver dysfunction, aspartate aminotransferase (AST) and alanine aminotransferase (ALT), were correlated with mortality. Although COVID-19 most commonly involves the respiratory system, these findings are consistent with clinical reports of severe disease from a generalized inflammatory state resulting in multiorgan damage and failure.

A subset of patients infected with SARS-CoV-2 experience prolonged recovery periods. In fact, in our multiscale embedding of patient clinical states we see a transition between a region of high survival likelihood score and a region of high extended recovery likelihood score (Figure 7A). In order to understand which cellular populations and clinical features drive the difference between these outcomes, we decided to zoom into this transition point (Figure 7B). We computed DREMI association scores between clinical features and flow sorted cellular populations to identify features differentially associated with survival and extended recovery. Our analysis found that age and kidney dysfunction were strongly associated with extended recovery indicating that older patients with worse kidney function were more likely to experience lengthy recovery periods from infection (Figure 7C).

Beyond clinical features like age and organ function, we also discovered different cellular populations associated with outcomes such as survival or long-term recovery from disease (Figure 7C), thus showcasing the way in which MultiScale PHATE can be used as a substrate for integrating and analyzing multiple modalities of data. Multiple recent publications have addressed the question of protection conferred by T cell-mediated responses [77–79] - with many demonstrating that antigen-specificity and T cell responsiveness lead to improved outcomes [80, 81]. Our analyses are in line with these findings, indicating that though some T cell subsets may be pathogenic, the major T cell subsets overall are positively associated with survival and negatively associated with a lengthy recovery phase. Interestingly, no myeloid subsets were found to be associated with length recovery periods, indicating that T cells and T cell subsets are perhaps more associated with recovery length while other immune populations may be associated with mortality.

## 5 Discussion

Here we presented a multiscale data exploration technique to visualize, understand and compare large-scale datasets, filling a key gap in biological data exploration. Multiscale PHATE finds groupings of data at varying scales that are predictive of clinical outcome. Biological data naturally contains multi-granular structure. Most analysis methods, however, whether clustering or dimensionality reduction algorithms, generally only look at a single level of resolution and do not offer a systematic way to explore different scales. Hierarchical clustering is one method that could offer certain scales of resolution. However, due to the constant merges that occur in hierarchical clustering approaches, like Louvain, many levels of resolution are missed and biologically important levels of granularity are not recapitulated. By contrast, Multiscale PHATE offers a fast manifold learning-based technique for uncovering a continuum of resolutions of structure and features from data. The speed and effectiveness of Multiscale PHATE is due in large part to key algorithmic advances we presented to make the underlying diffusion condensation process scalable, and representational advances in using diffusion potential coordinates as the substrate for the condensation.

We show that Multiscale PHATE can be combined with other techniques, like manifold density estimation (MELD), mutual information (DREMI), trajectory inference (TrajectoryNet), and classification (random forest classification) to provide deep and detailed insights in biological processes. We showed several effective examples of combining Multiscale PHATE with a technique known as MELD that shows the relative likelihood of seeing cells from different categories of patients in different parts of the cellular manifold. When MELD is combined with Multiscale PHATE, we can find levels of resolution that naturally capture the salient differences between patients with different clinical outcomes. Interestingly, Multiscale PHATE’s ability to zoom in helps identify pathogenic subsets of protective populations. Across our analyses, T cells have been shown to be protective against poor outcomes, corroborating previous work done in COVID-19. While broadly this cell type is protective, a multiscale zoom in of CD4^+^ T cells reveals a pathogenic CD4^+^IFNγ^+^GranzymeB^+^ Th17 subpopulation, highlighting the need for multiresolution analyses. In our work, we show several instances where density resampled estimate of mutual information (DREMI) between MELD likelihood scores and Multiscale PHATE identified clusters revealed potential associations between outcomes and key subpopulations, like CD16^+^CD14^−^ CD66b^−^ neutrophils and CD14^*lo*^CD16^+^HLA-DR^*lo*^ monocytes. We then showed that our manifold representation can be used to model cellular dynamics in patient response, identifying a key transition in Th17 cells as well as differential longitudinal IL-2 and IL-6 expression in patients with good and adverse outcomes. Furthermore, we showed that Multiscale PHATE identified populations combined with outcome variables can be used to predict clinical outcomes better than the current gold standard for flow cytometry analysis. Finally, we show that our approach is generalizable to a wide variety of biomedical data, including scRNAseq, scATACseq, CyTOF, TCR repertoire sequencing and clinical datasets.

While we have demonstrated Multiscale PHATE in the context of COVID-19 patient data, we believe that both the technique and the ways in which we have used it to analyze multi-modal data are widely applicable. Other applications could include analysis of multi-modal influenza or HIV data. Multiscale PHATE can also be used with an individual data modality to uncover structure where canonical cellular subtypes are not available, such as in patient-specific cancer or tumor cell types. Generally, as datasets continue to increase in size and the number of samples continue to expand, our scalable algorithm will become even more critical for joint analysis.

## 6 Acknowledgements

Yale IMPACT Research Team

Abeer Obaid^16^, Adam Moore^21^, Alice Lu-Culligan^3^, Allison Nelson^16^, Anderson Brito^10^, Angela Nunez^16^, Anjelica Martin^3^, Anne L Wyllie^9^, Annie Watkins^10^, Annsea Park^3^, Arvind Venkataraman^3^, Bertie Geng^16^, Chaney Kalinich^10^, Chantal BF Vogels^9^, Christina Harden^10^, Codruta Todeasa^16^, Cole Jensen^10^, Daniel Kim^3^, David McDonald^16^, Denise Shepard^16^, Edward Courchaine^17^, Elizabeth B. White^10^, Eric Song^3^, Erin Silva^16^, Eriko Kudo^3^, Giuseppe DeIuliis^12^, Haowei Wang^10^, Harold Rahming^16^, Hong-Jai Park^16^, Irene Matos^16^, Isabel M Ott^9^, Jessica Nouws^16^, Jordan Valdez^16^, Joseph Fauver^10^, Joseph Lim^18^, Kadi-Ann Rose^16^, Kelly Anastasio^19^, Kristina Brower^10^, Laura Glick^16^, Lokesh Sharma^16^, Lorenzo Sewanan^16^, Lynda Knaggs^16^, Maksym Minasyan^16^, Maria Batsu^16^, Maria Tokuyama^3^, M. Cate Muenker^16^, Mary Petrone^10^, Maxine Kuang^10^, Maura Nakahata^16^, Melissa Campbell^15^, Melissa Linehan^3^, Michael H. Askenase^20^, Michael Simonov^16^, Mikhail Smolgovsky^16^, Nathan D. Grubaugh^25^, Nicole Sonnert^3^, Nida Naushad^16^, Pavithra Vijayakumar^16^, Peiwen Lu ^3^, Rebecca Earnest^9^, Rick Martinello1^1^, Roy Herbst^16,23,24^, Rupak Datta^1^, Ryan Handoko^16^, Santos Bermejo^16^, Sarah Lapidus^9^, Sarah Prophet^16^, Sean Bickerton^17^, Sofia Velazquez^20^, Subhasis Mohanty^10^, Tara Alpert^1^, Tyler Rice^3^, Wade Schulz^22^, William Khoury-Hanold^3^, Xiaohua Peng^16^, Yexin Yang^3^, Yiyun Cao^3^ & Yvette Strong^16^

^16^Yale University School of Medicine, New Haven, CT, USA. ^17^Department of Biochemistry and Molecular Biology, Yale University School of Medicine, New Haven, CT, USA. ^18^ Yale Viral Hepatitis Program, Yale University School of Medicine, New Haven, CT, USA. ^19^Yale Center for Clinical Investigation, Yale University School of Medicine, New Haven, CT, USA. ^20^Department of Neurology, Yale University School of Medicine, New Haven, CT, USA. ^21^Yale University School of Public Health, New Haven, CT, USA. ^22^Center of Biomedical Data Science, Yale University, New Haven, CT, USA. ^23^ Yale Cancer Center, Yale New Haven Hospital, CT, USA. ^24^Smilow Cancer Hospital, Yale New Haven Hospital, New Haven, CT, USA. ^25^Department of Epidemiology of Microbial Diseases, Yale University School of Public Health, New Haven, CT, USA.

## Declaration of Interests

Dr. Krishnaswamy is on the scientific advisory board of KovaDx and AI Therapeutics.

Dr. Iwasaki a member of the SAB for InProTher.

Dr. Iwasaki is a co-founder of RIGImmune.

Dr. Wilson is founder of Efference.

Dr. Ko is a member of the expert panel of the Reckit Global Hygiene Institute.

## 7 Methods

### 7.1 Computational Methods

In the following sections we provide a thorough description of each aspect of the Multiscale PHATE algorithm and the use of downstream analysis tools. This includes but is not limited to explanations of algorithm design choices, information on how comparisons between algorithms were run and details on how the patient manifold was constructed.

#### 7.1.1 Diffusion information geometry for visualization and condensation

The multiresolution visualization provided by Multiscale PHATE relies on the construction of a diffusion geometry that captures the intrinsic structure of the data. Such a construction was first presented in the context of manifold learning with Diffusion Maps (DM), which rely on diffusion coordinates derived from spectral decomposition of the heat kernel over (Riemannian) manifolds [82]. The DM construction approximates the heat kernel on data by defining a Markovian diffusion process whose transition probabilities are given by 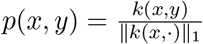, where the *L*_1_ norm is taken over the input data and *k*(·, ·) is a similarity kernel capturing local neighborhoods in the data. Then, an integral diffusion operator is constructed as 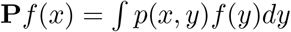, which is represented in finite settings as a matrix with entries [**P**]_*ij*_ = *p*(*x_i_, x_j_*), where {*x*_1_, *x*_2_,…} are the input data points (e.g., cells or strains in our case). By taking powers of this diffusion operator, we can consider *t*-step diffusion probabilities between data points given by 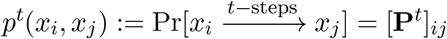. Finally, the diffusion geometry considers each data point *x* via its *t*-step diffusion distribution 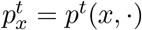, and DM aims to extract low dimensional coordinates where Euclidean distances capture a diffusion distance metric defined as *L*_2_ distances between these distributions, called *diffusion distance*.

While several similarity kernels are used in practice to construct the diffusion operator **P**, a standard choice is the Gaussian affinity *k_ε_*(*x, y*) = exp(−||*x* − *y*||^2^/*ε*) [13, 82–84], in which case we denote the diffusion operator **P**_*ε*_, where *ε* determines the neighborhood radius. This kernel choice is often seen in theoretical and mathematical work due to its established properties on data sampled from locally low dimensional geometries (i.e., data manifolds) [82, 85, 86]. In particular, it can be verified that when the data is sampled from a Riemannian manifold, the diffusion operator 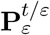 constructed from *k_ε_*(·, ·) converges to the heat kernel on the underlying manifold as *ε* → 0. Further, as *ε* → 0 the eigenvectors of 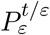 operator converge to Laplace-Beltrami eigenvectors that characterize the solutions of the heat equation 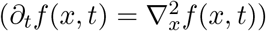 with Neumann boundary conditions on the underlying manifold of the data. Based on this convergence properties, the embedding provided by DM is based on the eigendecomposition of **P**^*t*^ to its eigenvalues 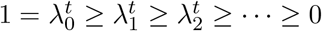 and corresponding eigenvectors *ϕ*_0_, *ϕ*_1_, *ϕ*_2_,…, which then yield the diffusion coordinates 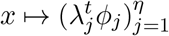, where *η* is determined by the numerical rank of **P**^*t*^ and the first eigenpair is discarded since φ_0_ is (provably) constant. We refer the reader to [82] for more details on DM and its properties.

While the analytic relation between spectral embedding with diffusion coordinates is appealing from a manifold learning perspective, the resulting DM often separates trajectories, pathways, or clusters into independent eigenspaces. This, in turn, yields multidimensional representations that cannot be conveniently visualized (e.g., having significantly more than 2-3 dimensions), and more importantly, cannot be directly projected into 2D or 3D displays that faithfully capture diffusion distances. In order to overcome this and extract a low dimensional data visualization, the recently proposed PHATE method treats the constructed diffusion geometry as a statistical manifold and leverages tools from information geometry to define a family of diffusion information distances defined as 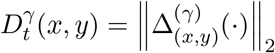 where

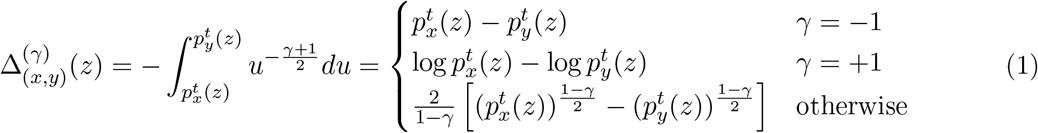

and the parameter −1 ≤ *γ* ≤ +1 attenuates the influence of lower probability differences in the overall distance. On one extreme (*γ* = −1), the resulting metric yields the traditional diffusion distance. When *γ* = 0, it yields an f-divergence corresponding to Hellinger distances between diffusion distributions. On the other extreme (*γ* = +1), the resulting information distance yields an *L*_2_ distance between localized diffusion energy potentials given by 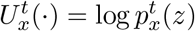, as discussed in [13]. There, as well as in other work [87], it has been shown that this potential distance is amenable to low dimensional embedding that captures and visually accentuates emergent global and local structures in the data - most importantly, trajectories and transitions between stable clusters in it. Therefore, the PHATE method, as well as its variation here, are based on embedding potential distances directly into two or three dimensional coordinates via stress-minimizing optimization procedure provided by multidimensional scaling (MDS). In addition to the core utilization of diffusion information geometry, the PHATE algorithm also includes robust construction of the initial neighborhood kernel, automatic tuning of diffusion resolution (see also discussion in the next section), and efficient sampling for scalability purposes. For more details about these aspects of PHATE, we refer the reader to [13].

Multiscale PHATE not only uses PHATE for visualization of several chosen iterations of the condensation process (explained below), representing multiple scales of data coarse graining, but also as the coordinate system for the data. This ensures that the condensation process itself operates in data manifold dimensions.

#### 7.1.2 Multiresolution via the diffusion condensation time-inhomogeneous Markov process

The diffusion geometry underlying PHATE is naturally multiscale, via the diffusion time parameter *t* that controls the resolution of information captured by the diffusion process. Indeed, as the diffusion time increases, the distributions 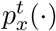 (or potentials 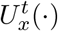) consider increasingly diffused energy that attenuates local differences until eventually at *t* → ∞ all these distributions converge to a unique equilibrium stationary distribution, since the process is ergodic. However, as discussed in [88, 89], this process often diffuses information too rapidly to enable multiresolution representation of varying intermediate scales of data geometry. Further, in [13], it was shown that the diffusion time scale admits an optimal time scale for visualization, which can be identified automatically by distinguishing between a rapid denoising phase and a slow decay from metastable to equilibrium diffusion states. We can use this property to automatically tune the diffusion time scale to transitioning point between these two phases. However, here we aim to extend this analysis to provide a full multiscale or multiresolution data geometry, and therefore we need to provide better control of the propagation of information by intrinsic diffusion over the data.

One of the first attempts at alleviating the rapid convergence to stationary distribution in multiscale DM was presented in [88], as part of a hierarchical construction of localized diffusion folders (LDF) using a localized diffusion process, which was further analyzed in [89]. The localized diffusion process there limited each instantiation of the diffusion random walks to only traverse between two “diffusion folders” (i.e., clusters), thus blocking global pathways that quickly diffuse to wide regions in the data. While this process was shown to be effective in some applications involving hierarchical clustering, it requires separate clustering steps and a priori determination of scales at which to pause the diffusion and cluster into LDFs. Furthermore, the pruning of the diffusion process there is computationally intensive, as each diffusion affinity (or transition probability) requires simulating or approximating a local diffusion process between two considered clusters. However, the principles posed by this approach clearly established the need for careful manipulation of the underlying Markov process of DM in order to truly enable multiscale representation learning via diffusion geometry, and by extension the diffusion information geometry used in PHATE.

A more recent approach towards multiresolution diffusion-based coarse graining was presented in [12], which relies on replacing the traditional time-homogeneous Markov process typically used in diffusion frameworks [13, 82] with an inhomogeneous process, following the theoretical analysis in [90] that established the mathematical viability of diffusion geometry construction of such processes. Unlike previous approaches, the coarse graining in [12] does not rely on a clustering & pruning approach. Instead, it proposes to base the intuition for the diffusion construction from heat propagation that rapidly spreads over the data based on connectivity, to a condensation process that alternates between slow gravitation (e.g., as drops of water slowly gravitate towards each other) and fast merging, concentrated regions collapse (e.g., as water drops merge together) to a single point. The alternation between these meta-stable and transient regimes also provides a diffusion-analogous notion of persistence used in topological data analysis (e.g., in the construction of persistence homology), which in turn naturally gives rise to emergent stable resolutions for multiscale visualization.

#### 7.1.3 Condensation on potential coordinates

The computation of the diffusion condensation process in [12] only uses the diffusion operator **P**, interpreted as a low-pass (smoothing) filter that can be applied to any dataset encoded in a points-by-features data matrix **X**. In the current manuscript rather than using the original features, we use the potential representation of data points used in PHATE (see Equation 1) as the as initial features. This effectively re-represents points by features that consist of the log of diffusion probabilities to all other features. In PHATE this representation is used as the step before dimensionality reduction and low-dimensional data embedding. However, we use these diffusion potential coordinates here as a high-dimensional representation of the data on which the condensation operates. This way when data points are condensed, they are condensed in terms of their diffusion probabilities.

Then, condensation process proceeds as follows. Let **X**^(0)^ = **X** be the initial data matrix with diffusion operator **P**_0_:= **P** constructed from its rows (as data points), and let **X**^(1)^ = **P**_0_**X**^(0)^. This gives the first iteration of the process, where the application of the diffusion smoothing intrinsically denoises and reduces local variability in **X**^(1)^ compared to **X**^(0)^. Then, the process is repeated to further reduce local data variability by computing the diffusion operator **P**_1_ over rows of **X**^(1)^, yielding **X**^(2)^ = **P**_1_**X**^(1)^. In general, this process is repeated iteratively, resulting in a time-inhomogeneous Markov process

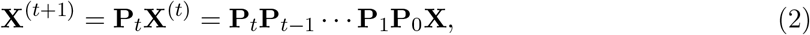

whose *t*-step transition probabilities are given by the entries of a time-varying row-stochastic operator **P**^(*t*)^ = **P**_*t*_⋯ **P**_0_. As mentioned previously, due to the low-pass nature of each diffusion operator **P**_*t*_, this Markov process adaptively removes local (high frequency) variations in input coordinate functions. The effect on the data points **X** is to draw them towards local barycenters, which are defined by the inhomogeneous diffusion process.

#### 7.1.4 Data merging

When two or more points collapse into the same barycenter (closer than a threshold ϵ), we merge them into a cluster since they would then have approximately the same coordinates. After this merging operation, we effectively treat the cluster as a single point. This has the effect of density subsampling the data iteratively, and allowing for subsequent iterations to proceed faster.

As we iterate this process over and over again, the condensation process slowly coarse grains the data to reveal structure at all levels of granularity while avoiding the typical tendency of traditional hierarchical clustering approaches to force (e.g., greedy) cluster merges at every scale.

#### 7.1.5 Distinction and comparison between the diffusion condensation process and hierarchical clustering

One use of diffusion condensation can be to provide a hierarchy of clusters determined by merged points. However, it should be noted that the condensation process here is significantly different from typical hierarchical clustering, and instead provides a richer coarse graining of data geometry. Indeed, hierarchical clustering algorithms generally belong to two families: divisive algorithms and agglomerative ones.

Divisive approaches (e.g., bisecting *k*-means [91] or MST-based clustering [92]) work in a top-down fashion, each time optimizing a partition of the data into clusters (e.g., using partitional methods like *k*-means), and then recursively partitioning further each cluster. The difference between these and the gradual aggregation approach of the condensation process is clear.

Agglomerative methods, on the other hand, work in a bottom-up fashion by first merging points into clusters, and then recursively merging increasingly bigger clusters. While intuitively more related to the gradual merges in diffusion condensation, there is a fundamental difference between the coarse graining operation applied here and the (typically greedy) agglomeration in such methods. Indeed, most agglomerate clustering methods only operate on determining an iterative or recursive sequence of merges, without considering any intermediate information or structure in the data.The condensation process utilized here, on the other hand, is derived from a continuous process that gradually eliminates local variability in the data. At its core, it relies on a time-inhomogeneous Markov chain that gradually constructs a diffusion geometry that reveals global and local structures in the data at increasingly coarse scales. The elimination of local variability in this process allows points to naturally come together, thus producing natural data clusters from data regions that collapse to the same point, without the need for partitioning or greedy agglomeration. However, this is a pattern that emerges from the coarse graining process, rather than directly or explicitly guiding it. The constructed multiresolution data geometry also reveals other information, beyond clustering, which makes it amenable for visualization and other downstream tasks. For instance, condensation tree produces branch lengths that are meaningful, and levels of meta-stability can be analyzed, as we do for the selection of meta-stable resolutions (e.g., for visualization) explained below.

To demonstrate the difference between diffusion condensation and agglomerative clustering, we use the Louvain method [28] as a representative example, due to its popularity in single cell data analysis. This method greedily selects clusters to merge together by their impact on modularity (i.e., whether and how much they improve it). The forced merges, while ensuring a hierarchy of data agglomerations, do not provide reliable coarse grained representations for revealing varied data resolutions. As we showed in figure 3 and supplementary Figure 3 miss vital levels of resolution. Meanwhile, diffusion condensation allows for a systematic exploration of granularity and is better at capturing levels where biological differences may exist (Supplementary Figure 3B).

In order to quantitatively compare the accuracy of Multiscale PHATE clusters with hierarchical clustering approaches, we compared cluster labels generated from a range of clustering strategies to ground truth labels using Adjusted Rand index (ARI). We first generated a hierarchical stochastic block model with different clusters at multiple granularities (Figure 3A, Supplementary Figure 3A). We then used Multiscale PHATE, Louvain [28], Leiden [29] and single linkage hierarchical clustering [30] to identify groupings across multiple levels of granularity. For each level of ground truth clusters, we computed ARI against cluster labels from each algorithm across all granularities, storing the highest ARI for each method. For the flow cytometry data, we used gated populations from 3 samples in our myeloid-centric flow cytometry panel as ground truth labels across coarse and fine grain cluster labels. For instance, at coarse grain monocytes would be identified as one population, however at fine grain monocytes would be a part of three distinct populations. ARI was computed similarly for this dataset, ground truth labels were compared to all granulities of clusters from each algorithm, with the top score stored for each approach. Networkx [93] was used to produce Louvain clusters, leidenalg was used to produce Leiden clusters and agglomerative clusters were produced using sklearn [94].

#### 7.1.6 Scalable coarse-graining with fast diffusion condensation

In order to allow Multiscale PHATE to enable scalable exploration of large data sets, such as high dimensional biological data, we propose speeding up of the initial condensation iteration in the following ways:

1. Speed-up of the initial iteration using graph partitioning.
2. Fast computation of the diffusion potential via landmarking.

The complexity of computing a diffusion operator on *n* points is *n*^2^. However as the condensation proceeds, points that are within a threshold distance of each other are merged into a single point. Therefore the number of points steadily decreases, allowing the algorithm to speed up in successive iterations. This process means, however, that the first calculation of the operator is the most computationally complex. In order to reduce the dimensionality of the initial condensation iterations, we run hierarchical kmeans on the PCA space of the data with a high K (by deafult 100) to obtain a coarse graining of the data in feature space. In each iteration of the kmeans approach we partition the data into k more clusters. In following iterations we compute another k clusters from each of these clusters. This process continues until we have a large number of clusters from which to compute the diffusion operator (by default 25,000). We then compute a landmarked diffusion potential (as done in [13] and explained below) on this reduced space, by convention the centroid of each of these clusters, before starting the coarse graining process.

Creation of the diffusion operator requires the computation of all pairwise distances between points, before conversion of those distances to affinities. Instead using spectral clustering on a set of data we can come up with cluster centroids that are treated as “landmarks”, Distances **D**_*pl*_ and **A**_*pl*_ are computed between points and landmarks, i.e., they are *n* × *k* matrices where *n* is the number of points and *k* is the number of landmarks. In addition, distances *D_l_* and affinities **A**_*l*_ are computed between landmarks, i.e. *k* × *k* matrices. Then in order to compute the diffusion operator **P**^*t*^ we row normalize **A**_*pl*_ and **A**_*l*_ to obtain **P**_*pl*_, **P**_*l*_ and compute 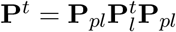 decomposing *t*-step path probabilities between two points as the probability of going to a landmark and then back to the point. We have shown in [13] that this leads to high quality approximations of the diffusion operator which lead to near-identical visualizations with PHATE. In addition, we examined in [19] that this leads to low error approximations of diffusion operators in general. We use this fast approach to compute a low error diffusion potential system for our coarse graining process.

We show that the resultant method is orders of magnitude faster than competing methods: DM, tSNE, UMAP, Monocle2, and PHATE (Figure 2B).

#### 7.1.7 Selection of visualization layers via Gradient Analysis

As our iterative coarse graining approach creates hundreds of layers for downstream analysis, selecting salient level of granularity for visualization is a critical task. As previously described, Multiscale PHATE visualizes emergent stable resolutions that define the information geometry of the manifold well at a particular scale. In order to identify these metastable states, we identify changes in manifold density in for every pair of successive condensation iterations ∇^(*t,t*+1)^ = **X**^(*t*)^ – **X**^(*t*+1)^. In order to identify total shifts in density we compute the matrix sum of ∇^(*t,t*+1)^ by 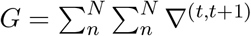. Picking scales for visualization and downstream analysis arises from identifying local minima in *G*.

#### 7.1.8 Comparison of hierarchical visualizations with DeMAP

DeMAP is a metric for assessing visualization and dimensionality reduction quality in terms of its ability to capture the manifold geometry of the data even when the data has noise and perturbations. It was first proposed in [13]. DeMAP computes correlation between geodesic distances on ground truth noiseless data manifolds to Euclidean distances in the embedded space, after adding noise to the data. Hence it is called *Denoised Manifold Affinity Preservation*. High DeMAP scores indicate that a visualization that accurately represents geodesic manifold distances in an embedding.

In order to show that Multiscale PHATE created improved hierarchical visualizations when compared to other approaches, we performed an ablation study.

First, the *splatter* software was used to simulate ground truth and noisy single cell data of either group (cluster) or path (trajectory) geometries [26]. After computing the condensation tree on the noisy splatter data, we created a Multiscale PHATE embedding by identifying the optimal resolution via gradient analysis. In order to create hierarchical visualizations of other algorithms, we selected the same resolution and merged points together in raw feature space to create a dataset of the same scale across both noisy and ground truth datasets. After computing other dimensionality reduction techniques on the merged noisy data, we compared all embeddings with the merged ground truth data using DeMAP. This process was repeated across a range of noise types, biological variation and drop out, and a range of noise levels. For robustness, this process was run across 10 different splatter datasets with group geometry and 10 different splatter datasets with path geometry. Besides Multiscale PHATE, the DeMAP package was used to build all visualizations [13].

#### 7.1.9 Construction of patient manifold through multiresolution cluster evaluation

Previously, PhEMD built a manifold of samples measured via single cell technologies by binning cells associated with each sample into histograms and computing Earth Mover Distances or optimal transport between histograms [22]. This computation, however, is done at a single scale and requires determining ground distance between histogram bins. Truthfully, cells can occupy a diverse set of hierarchical labels which a single resolution does not capture. Here, instead, we combine multiple histograms at different scales for each patient to compute a distance between their underlying cellular states. Our approach of replacing ground distance with multiscale construction is based on the work of [24] and [23], which show that with appropriate weights, the combination of such smoothed data distributions can be used to efficiently compute or approximate Earth mover distances. We note that our results show empirically that even without careful tuning of such weights, the resulting patient to patient distance, and the constructed manifold, accurately recapitulate the clinical states.

Practically, we create a manifold of samples by simultaneously evaluating multiple levels of the diffusion condensation tree. At each level *ℓ* ∈ {1,2,…, *L*}, a number of *N_ℓ_* clusters are identified. We count the number of cells, *n_ℓ,j,k_*, of the *k*-th patient that belong to each cluster *C_ℓ,j_* for every *j* ∈ {1, 2,…, *N_ℓ_*} and calculate the normalized percentage as 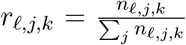. We calculate the proportions for all patients at a series of selected levels of the tree and concatenate these to create a rich multiscale vector of features for each patient. These multiscale feature vectors are then used to create an embedding with PHATE [13] and to de-noise patient specific signals using MAGIC [18] using Euclidean distance between samples.

#### 7.1.10 Use of MELD with Multiscale PHATE

MELD is a method proposed in [14] that takes a discrete signal defined on a data graph and computes a continuous likelihood score of the signal value by using a sophisticated form of neighborhood averaging by using a heat kernel at each point (Figure 1C). In order to apply MELD to this dataset, we combined the flow cytometry data coming from all patients, and used a binary outcome score that we call *mortality*, which uses a discrete 0-value for a positive outcome (the patient was discharged), or a 1-value for a negative outcome (patient died or was sent to hospice). The outcome of the patient is used as the discrete condition for all cells from that patient. Thus in our combined flow-cytometry dataset, every cell from positive outcome patients get a raw experimental signal value of 0. Using MELD, we estimate likelihood of each outcome over the cellular manifold using a heat-diffusion kernel applied to the data graph to obtain mortality likelihood score. Values of the mortality likelihood score range from 0 to 1 and constitute a probability likelihood estimate of the condition over the manifold. This allows us to identify areas of the cellular manifold that are likely to be enriched in those with positive or negative outcomes.

Since Multiscale PHATE identifies clusters of cells across all levels of granularity, we can sweep across resolutions to identify levels which isolate high and low mortality likelihood score regions. In fact, when comparing our multigranular clusters with other clustering techniques across a range of granularities, we show that multiscale PHATE is better able to isolate high and low mortality likelihood score regions in one of our flow cytometry panels (Supplementary Figure 3B). By looking at these informative resolutions, we can identify populations of cells that are pertinent to patient outcomes. When identifying these subpopulations in conjunction with cell type defining markers, we show that we can identify cell types and functional subtypes that are differentially enriched across patient outcomes and may drive disease pathogenesis. The full Multiscale PHATE and MELD integrated pipeline can be seen in Supplementary Figure 1B.

#### 7.1.11 DREMI Associations with mortality likelihood score

DREMI or Density Resampled Estimate of Mutual Information [15], is an information-theoretic metric that quantifies associations or strength of a relationship between two variables. Like most discrete estimates of mutual information, DREMI starts by binning continuous data into equal-sized partitions, *X* = {*X*_1_, *X*_2_,… *X_n_*}, and *Y* = {*Y*_1_, *Y*_2_,… *Y_n_*} in both variable dimensions but instead of measuring the mutual information as *I*(*X, y*) = *H*(*Y*) – ∑_*i*_ *H*(*Y*|*X_i_*) the difference between the entropy of *Y* and the conditional entropy of *X|Y*, DREMI “resamples” or equalizes the number of samples in each bin using an extra level of conditioning. Thus DREMI computes *DREMI*(*X, Y*) = *I*(*X, Y|X*) = *H*(*Y|X*) – ∑_*i*_ *H*(*Y|X_i_|X_i_*). The rationale for this is that normal mutual information is dominated by the density peaks of the *X* variable, and does not reveal the full strength of the relationship given imbalanced sampling which is common in biomedical data.

When combining our DREMI analysis with previously computed mortality likelihood score, we can identify functional marker trends which are correlated with mortality. As cells of the same type can occupy a range of functional states that can be enriched in disease, a given subtype may not be associated with mortality but a functional substate could be. By computing DREMI associations between mortality likelihood score and cellular functional state markers, we can identify markers, and by extension activation states, that are associated with outcome.

#### 7.1.12 Trajectory analysis of flow cytometry Data with TrajectoryNet

TrajectoryNet [21] trains a model to infer developmental and activation trajectories from a series of static snapshot measurements. The time-lapsed measurements are treated as samples from distributions at different time points. TrajectoryNet generates dynamic optimal transport between these distributions by restricting the paths taken using an path-derivative regularization, effectively interpolating between the time points and generating continuous differentiation paths [21].

TrajectoryNet builds upon work on Neural Ordinary Differential Equations (Neural ODE) [95], a new family of deep neural network models that learn a high dimensional derivative (rather than a function) with a neural network, and computes both the output, and backpropagates using an ordinary differential equation solver. More specifically, the ODE that describes the continuous dynamics of input variables 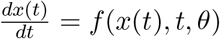, where *θ* are the neural network parameters. TrajectoryNet extends Neural ODE by adding regularization to the optimization target, which are specifically beneficial for modeling biological trajectories. We used the basic model of TrajectoryNet adopts an energy regularization in the form of 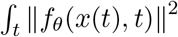, which allows for achieving optimal transport, especially suitable for cell differentiation trajectories.

We train TrajectoryNet to infer activation dynamics in our CD4^+^ T cell flow cytometry data by making use of clinical time series data from each patient sample. As each sample was taken some amount of time after presentation of initial infection symptoms, ranging from several hours to 60 day, we used this variable as explicit time. In order to run TrajectoryNet on these samples, we discretized the time variables into 4 different quartiles, representing timepoints 1 through 4 and trained TrajectoryNet to infer differentiation trajectories between these time points. To train TrajectoryNet, two types of data are used, the normalized flow cytometry data and the embedding from Multiscale PHATE. The embeddings are low-dimensional representations of the original data, and trajectories in the embedding space allow us to understand cellular dynamics over time, such as the differentiation of homeostatic Th17 cells into more pathogenic Th17 cells. On the other hand, trajectories in the original high-dimensional flow cytometry space allow for us to study time dependent changes in marker expression. This analysis allows for us to derive biomarkers, like IL-2 and IL-6, that are divergent and potentially predictive of patient outcome.

#### 7.1.13 Patient manifold analysis from Multiscale PHATE Features

In order to identify the differences between individual patient samples, we used Multiscale PHATE to construct a manifold of patients as described above. Similar to mortality likelihood score computed by MELD in our flow cytometry analysis, we computed a similar mortality likelihood score for our patient manifold by identifying if each patient sample originated from a patient that had a positive outcome or a negative outcome. In order to identify patient sample features correlated with mortality likelihood score, we compiled a set of clinical, demographic and Multiscale PHATE identified cell type proportion features for each patient sample. Using the geometry of the patient manifold, we de-noised our patient sample features using MAGIC [18] before running association analysis between features using DREMI [15].

#### 7.1.14 Mortality Prediction using Random Forest Classifier

In addition to being useful for visualizing, clustering and identifying condition specific enrichment of cell types, we wanted to see if the populations we identified across granularities were predictive of patient outcome. In order to predict patient outcomes from just a single patient sample, we trained a random forest classifier on populations we identified in our myeloid focused flow cytometry panel. Similar to our patient manifold analysis, we derived multiscale patient features by identifying the proportion of each patient’s cells that were labeled with a particular cell type. After partitioning our dataset of 210 patient samples into 5 sets, we performed 5-fold cross-validation where we iteratively shuffled training sets (4 of 5) and test sets (1 of 5). Across all runs, we achieved an accuracy of 83.5% across both conditions, with 86.5% classification accuracy for patients that survive and 78.8% classification accuracy for patients that died from infection. In order to identify cellular types that were particularly informative of mortality outcome, we computed and compared the feature importance of random forests. This analysis revealed that that monocytes, CD16^+^ neutrophils, dendritic cells and T cells had the highest importance and were most predictive of mortality outcome. To determine whether our Multiscale PHATE derived cellular populations were more informative than current gold standard cell typing strategies, we also trained a random forest classifier on cell populations identified via conventional flow cytometry gating analysis. This analysis was only able to accurately predict patient outcomes in 73.8% of cases.

#### 7.1.15 Software availability

The Multiscale PHATE package, as implemented in python, is available for download with a guided tutorial on the Krishnaswamy Lab Github page: https://github.com/KrishnaswamyLab/Multiscale_PHATE.

#### 7.1.16 Pre-processing of patient flow cytometry data

Flow cytometry was performed on PBMC from each patients (methods explained in detail below). The resulting .FCS files were pre-processed by applying compensation based on the respective single-color compensation controls, 2) selecting only leukocytes and singlets based on FSC and SSC, and 3) selecting only live cells based on a viability dye. MFI values for each fluorophore on a per-cell basis were then extracted for downstream analysis. In order to extract T cells for the cytokine focused T cell panel, cells with CD3 staining greater than 425 were extracted. For the T cell surface marker panel, cells with a CD3 staining greater than 500 were extracted. For the B cell focused panel, cells with a CD19 staining greater than 400 were extracted and cells expressing less than a total of 2700 cumulative staining across all markers were removed. No extraction of cells was done for the myeloid focused panel, however cells with cumulative staining across all markers less than 2700 across were removed. All datasets were then independently normalized to 1000 staining counts per cell before square root normalization.

### 7.2 Biological and Medical Methods

In the following sections we provide details on how patient biological data and clinical information was acquired and processed.

#### Ethics statement

This study was approved by Yale Human Research Protection Program Institutional Review Boards (FWA00002571, protocol ID 2000027690). Informed consent was obtained from all enrolled patients and healthcare workers.

#### Patients

Patient enrollment, sample acquisition, processing, and downstream analysis by flow cytometry were performed as in [25]. One-hundred and sixty-eight patients admitted to YNHH with SARS-CoV2 between 18 March 2020 and 27 May 2020 were recruited to the Yale IMPACT study (Implementing Medical and Public Health Action Against Coronavirus CT) after testing positive for SARS-CoV2 by qRT-PCR and included in this study. No statistical methods were used to predetermine sample size. Paired whole blood for flow cytometry analysis was collected simultaneously in sodium heparin-coated vacutainers and kept on gentle agitation until processing. All blood was processed on the day of collection. Patients were scored for COVID-19 disease severity through review of electronic medical records (EMR) at each longitudinal time point. For all patients, days from symptom onset were estimated as follows: (1) highest priority was given to explicit onset dates provided by patients; (2) next highest priority was given to the earliest reported symptom by a patient; and (3) in the absence of direct information regarding symptom onset, we estimated a date through manual assessment of the electronic medical record (EMRs) by an independent clinician. The clinical data were collected using EPIC EHR and REDCap 9.3.6 software. At the time of sample acquisition and processing, investigators were unaware of the patients’ conditions. Blood acquisition was performed and recorded by a separate team. Information about patients’ conditions was not available until after processing and analysis of raw data by flow cytometry and ELISA. A clinical team, separate from the experimental team, performed chart reviews to determine relevant statistics. Flow cytometry analyses were performed blinded. Patients’ clinical information and clinical score coding were revealed only after data collection.

#### Isolation of PBMCs

PBMCs were isolated from heparinized whole blood using Histopaque (Sigma-Aldrich, #10771-500ML) density gradient centrifugation in a biosafety level 2+ facility. After isolation of undiluted serum, blood was diluted 1:1 in room temperature PBS, layered over Histopaque in a SepMate tube (StemCell Technologies; #85460) and centrifuged for 10 min at 1,200g. The PBMC layer was isolated according to the manufacturer’s instructions. Cells were washed twice with PBS before counting. Pelleted cells were briefly treated with ACK lysis buffer for 2 min and then counted. Percentage viability was estimated using standard Trypan blue staining and an automated cell counter (Thermo-Fisher, #AMQAX1000).

#### Flow cytometry

In brief, freshly isolated PBMCs were plated at 1–2 × 106 cells per well in a 96-well U-bottom plate. Cells were resuspended in Live/Dead Fixable Aqua (ThermoFisher) for 20 min at 4 °C. Following a wash, cells were blocked with Human TruStain FcX (BioLegend) for 10 min at RT. Cocktails of desired staining antibodies were added directly to this mixture for 30 min at RT. For secondary stains, cells were first washed and supernatant aspirated; then to each cell pellet a cocktail of secondary markers was added for 30 min at 4 °C. Prior to analysis, cells were washed and resuspended in 100μL of 4% PFA for 30 min at 4 °C. For intracellular cytokine staining following stimulation, cells were resuspended in 200μL cRPMI (RPMI-1640 supplemented with 10% FBS, 2 mM l-glutamine, 100 U/ml penicillin, and 100 ug/ml streptomycin, 1 mM sodium pyruvate, and 50μM 2-mercaptoethanol) and stored at 4 °C overnight. Subsequently, these cells were washed and stimulated with 1× Cell Stimulation Cocktail (eBioscience) in 200 μL cRPMI for 1 h at 37 °C. 50μL of 5x Stimulation Cocktail (plus protein transport inhibitor) (eBioscience) was added for an additional 4 h of incubation at 37 °C. Following stimulation, cells were washed and resuspended in 100 μL of 4% PFA for 30 min at 4 °C. To quantify intracellular cytokines, these samples were permeabilized with 1× permeabilization buffer from the FOXP3/Transcription Factor Staining Buffer Set (eBioscience) for 10 min at 4 °C. All subsequent staining cocktails were made in this buffer. Permeabilized cells were then washed and resuspended in a cocktail containing Human TruStain FcX (BioLegend) for 10 min at 4 °C. Finally, intracellular staining cocktails were added directly to each sample for 1 h at 4 °C. Following this incubation, cells were washed and prepared for analysis on an Attune NXT (ThermoFisher). Data were analysed using FlowJo software version 10.6 software (Tree Star).

#### Acquisition of Clinical Data for Flow Cytometry analysis and Patient Manifold

Longitudinal patient data was extracted from the electronic medical record (Epic, Verona, WI) for only the hospitalized patients included in the repository. Time-varying data, specifically vital signs as well as laboratory studies, were extracted specifically 24 hours before and after the collection of blood specimens for flow cytometry as described above. This ensured that the measurements correlated with the patient state at the time of flow cytometry measurements. Laboratory values reflecting clinical evaluation of general inflammatory states (white blood cell count, high sensitivity c-reactive protein) were extracted. The values for the laboratory measurements were then consolidated by taking the most abnormal value (e.g. highest ferritin) in the 72 hour period and overlaid onto the patient manifolds.

#### Acquisition of Clinical Data for Clinical Manifold

For patients who did not undergo flow cytometry analysis, the time varying clinical, laboratory, and treatment data was extracted for the first 24 hours from admission with consolidation by the most abnormal value as described before. Otherwise, the consolidated data temporally correlating to flow cytometry measurements were extracted as described above.

## 8 Supplementary Figures

**Supplementary Figure 1:**
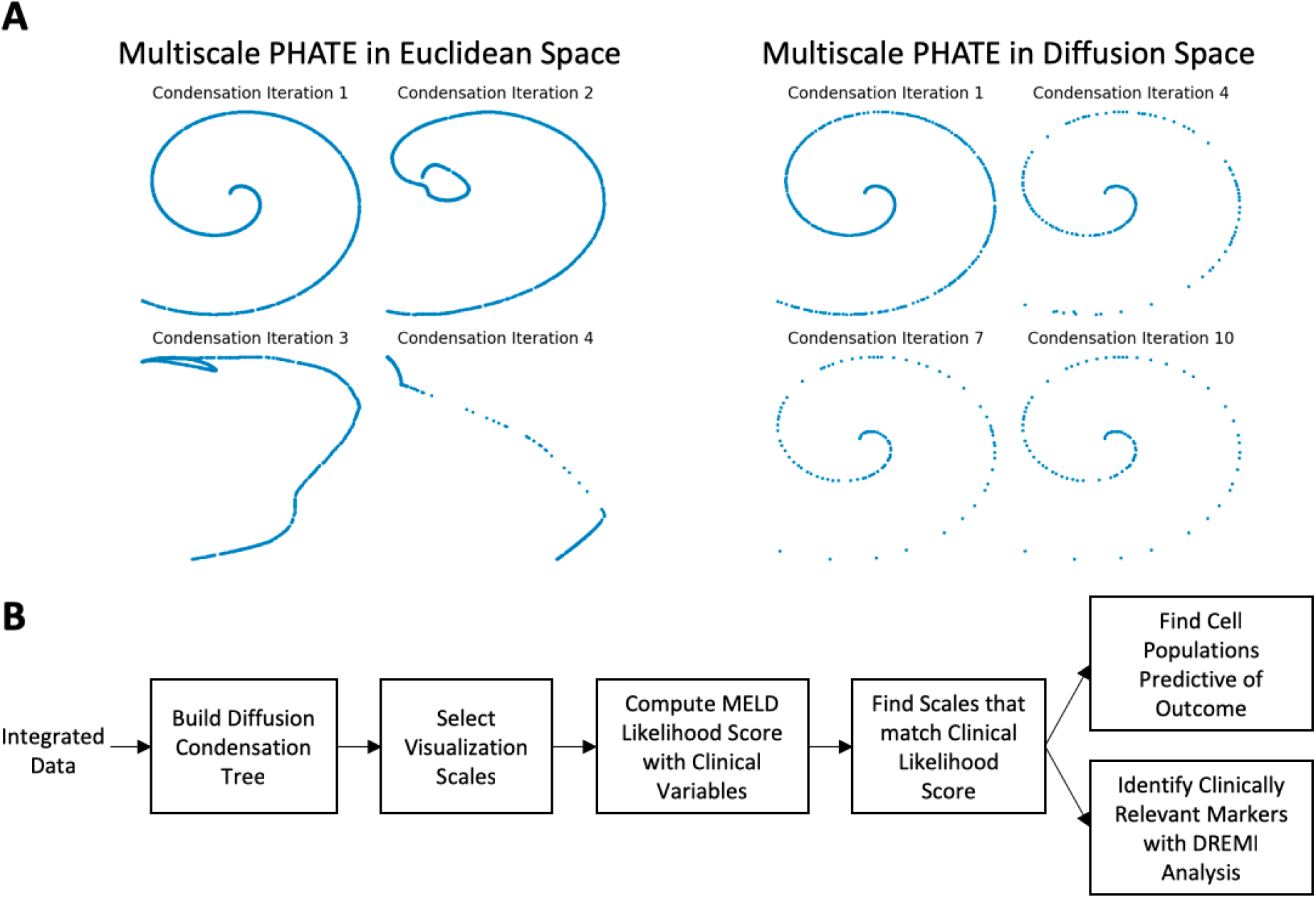
Condensing on Manifold and updating visualization. A. Visualization of toy swiss roll dataset after several iterations of fast diffusion condensation, running in both feature space and in manifold space as computed by Diffusion Potential. B. Pipeline for identifying cellular populations enriched based on clinical variables with Multiscale PHATE and MELD.

**Supplementary Figure 2:**
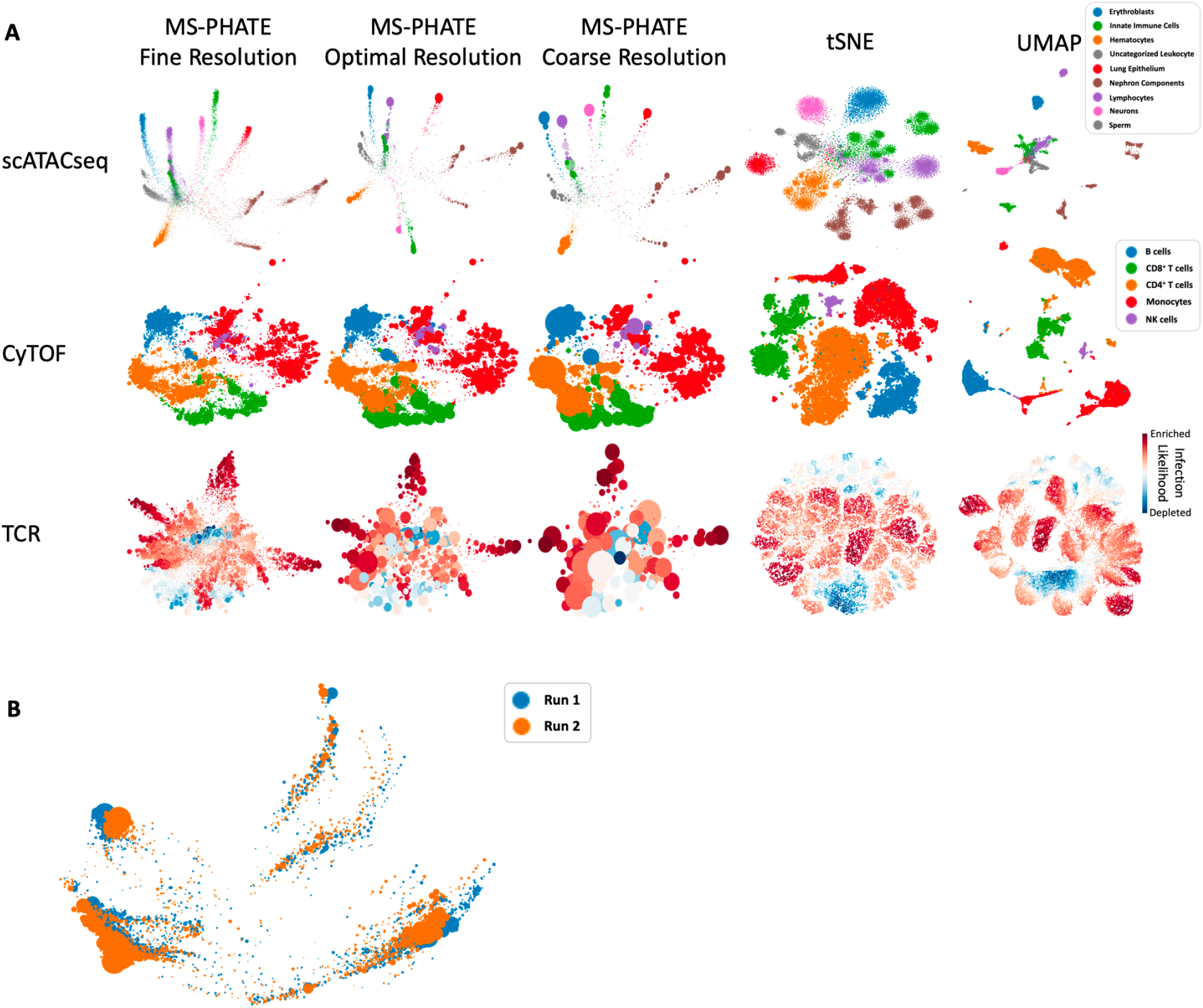
Visualization of additional data types with run time and reproducibility comparison. A. Visualization comparison of 25,528 cells from a diverse set of mouse tissues measured by scATACseq [96], 1,010,964 PBMCs measured by CyTOF [97] and 50,000 TCRs from COVID-19 infected patients and healthy controls [98, 99]. B. Visualization of reproduciblity of Multiscale PHATE across two different runs.

**Supplementary Figure 3:**
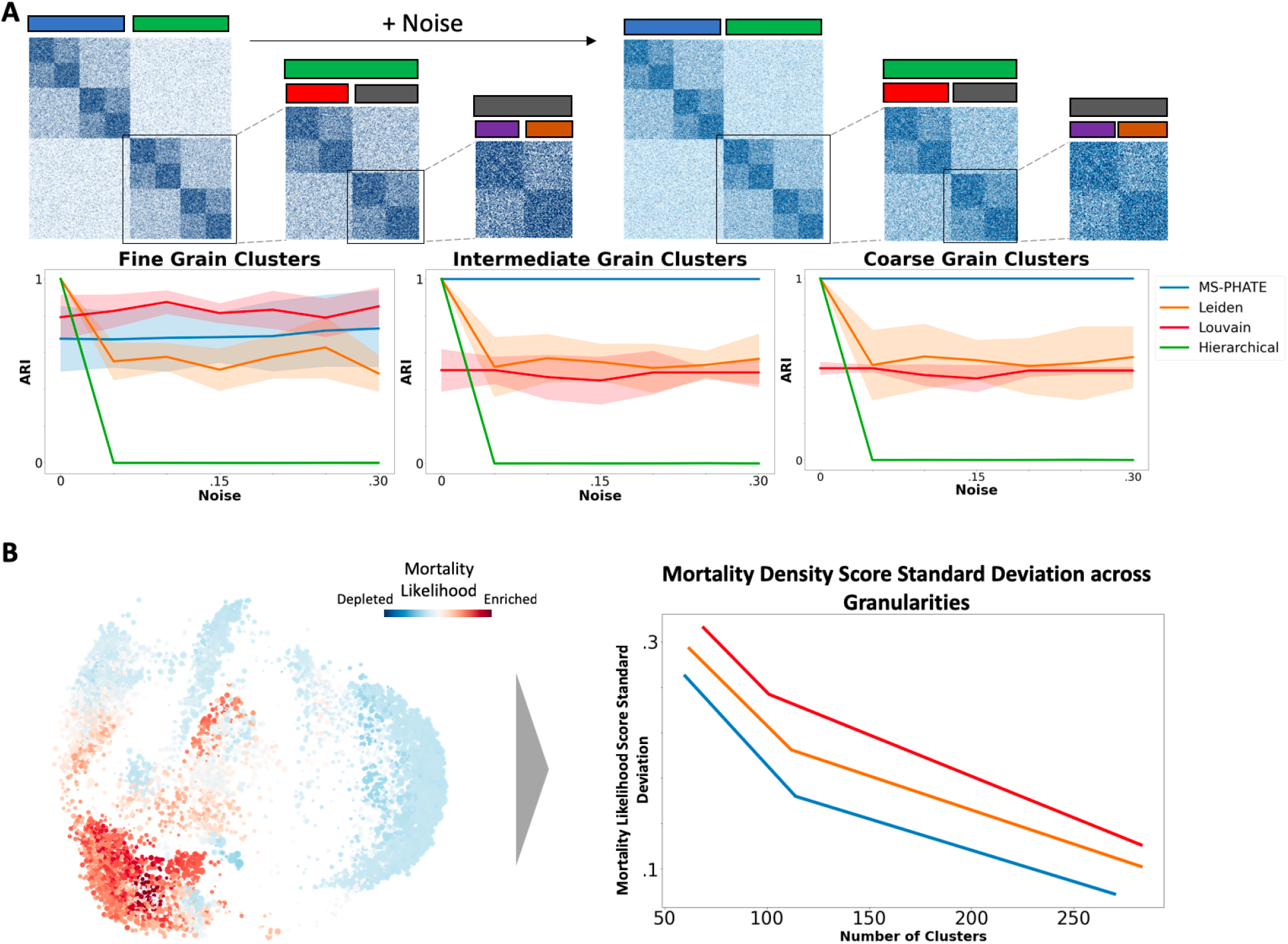
Comparison of Multiscale PHATE algorithm with other clustering techniques on three-layer hierarchical SBM and at identifying MELD density score enriched regions. A. Schematic of three-layer hierarchical SBM we generated for multigranular cluster comparison. For each method, increasing amounts of random gaussian values were added to the adjacency matrix of stochastic block model to simulate increasing amounts of noise. As the model was constructed with known clusters at multiple scales, we computed Adjusted Rand Index (ARI) between each algorithms predicted clusters and the known clusters across coarse and fine granularities. B. Comparison of multiple clustering techniques at identifying regions with uniform MELD likelihood scores across a range of comparable granularities.

**Supplementary Figure 4:**
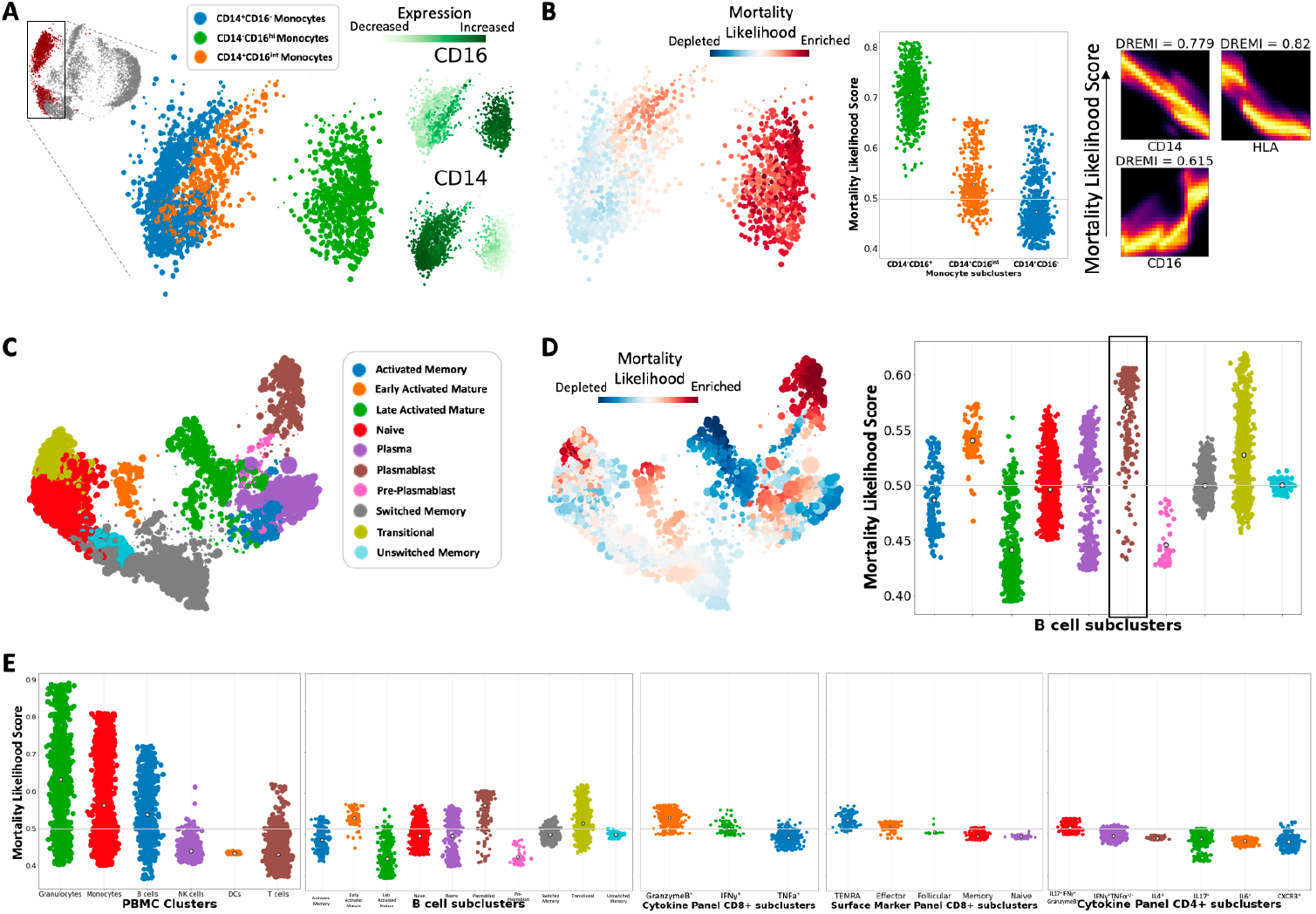
Multiscale PHATE identifies subsets of monocytes and B cells enriched in patients who die from COVID-19. A. Zoom in of monocyte population identifies subsets based on expression of markers. B. Visualization of mortality likelihood score in monocytes identifies subsets enriched in patients who die from COVID-19. Key associations between markers and mortality likelihood score computed by DREMI and visualized with DREVI. C. Visualization of B cells panel identifies a range of subsets based on expression of known markers. D. Visualization of mortality likelihood score identifies B cell subsets enriched in patients who die from COVID-19. E. Comparison of mortality likelihood score across panels reveals that granulocytes and monocytes are broadly the most enriched cell types in patients who die from COVID-19.

**Supplementary Figure 5:**
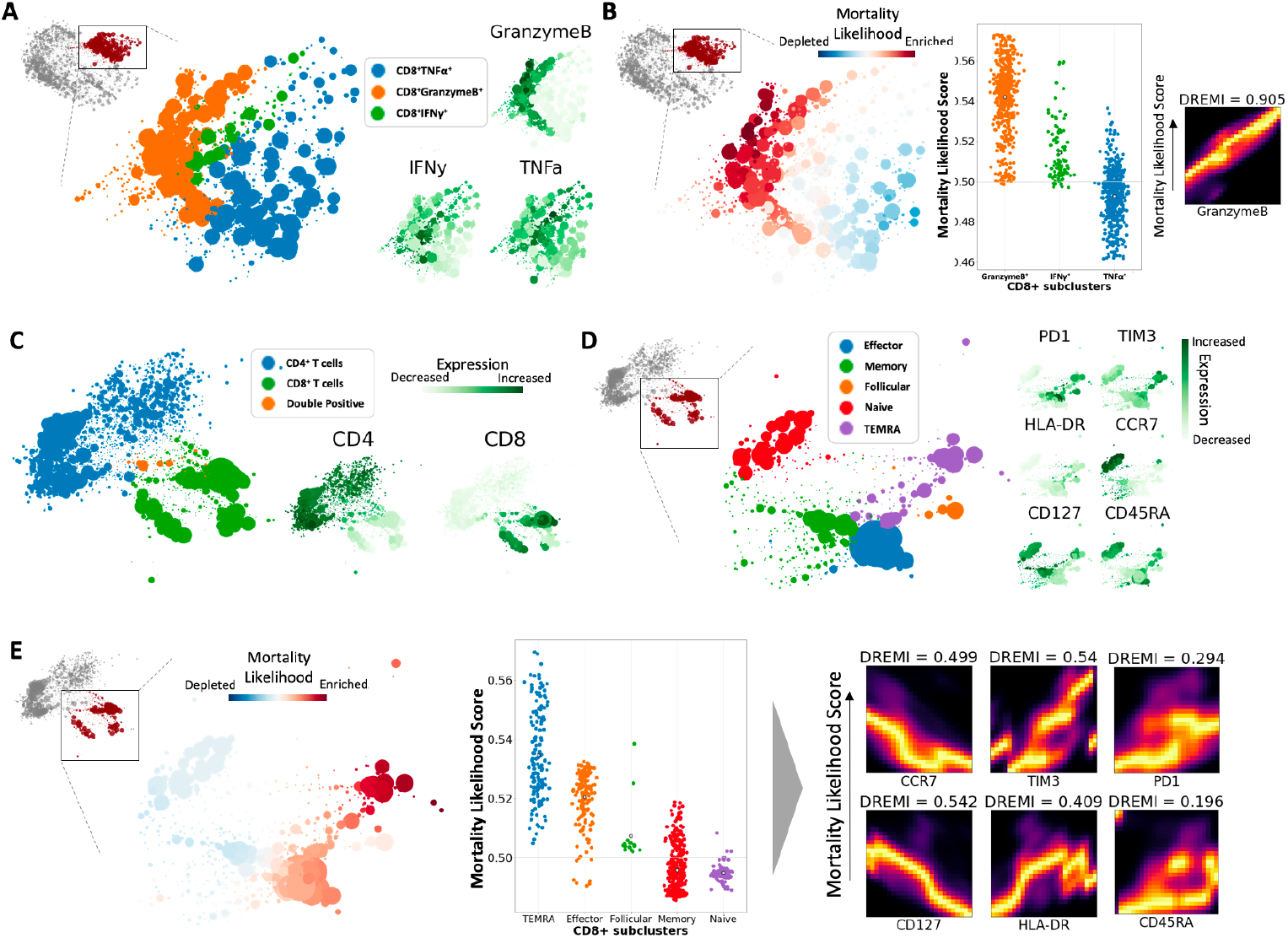
Multiscale PHATE analysis identifies subsets of CD8^+^ T cells enriched in patients with poor COVID-19 outcomes. A. Zoom in of CD8^+^ T cells identifies subsets based on expression of markers. B. Visualization of mortality likelihood score in CD8^+^ T cells identifies subsets enriched in patients who die from COVID-19. Key associations between GranzymeB and mortality likelihood computed by DREMI and visualized with DREVI. C. Multiscale PHATE visualization of T cell focused surface marker panel with broad T cell subtypes identified. D. Zoom in of CD8^+^ T cells identifies subsets based on expression of known markers. E. Visualization of mortality likelihood score in CD8^+^ T cells identifies subsets enriched in patients who die from COVID-19. Key associations between markers and mortality likelihood computed by DREMI and visualized with DREVI.

**Supplementary Figure 6:**
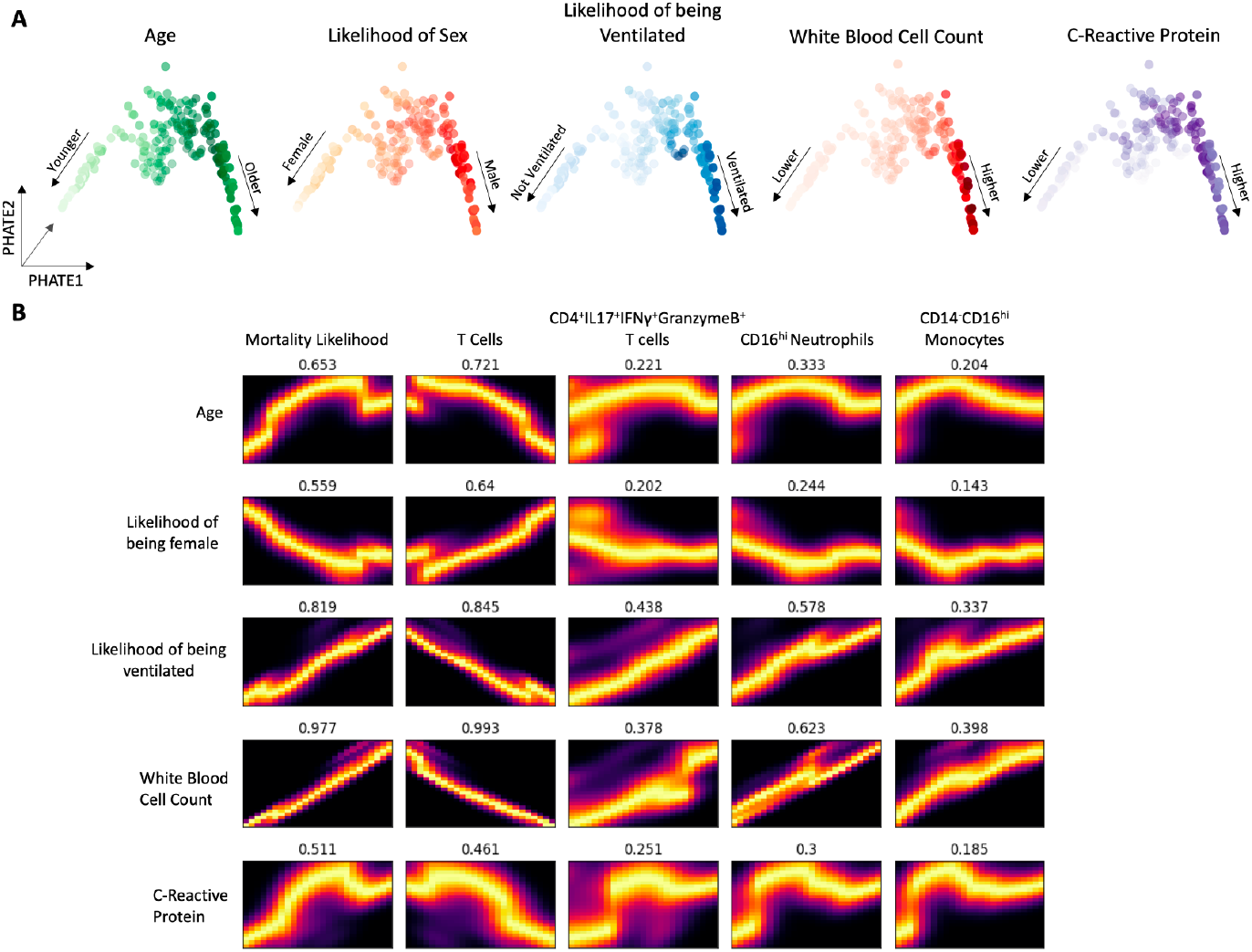
Visualization of patient manifold and correlation with clinical features. A. Visualizing clinical trends on patient manifold. B. DREMI and DREVI association analysis between clinical features and mortality as well as cellular populations.

**Supplementary Figure 7:**
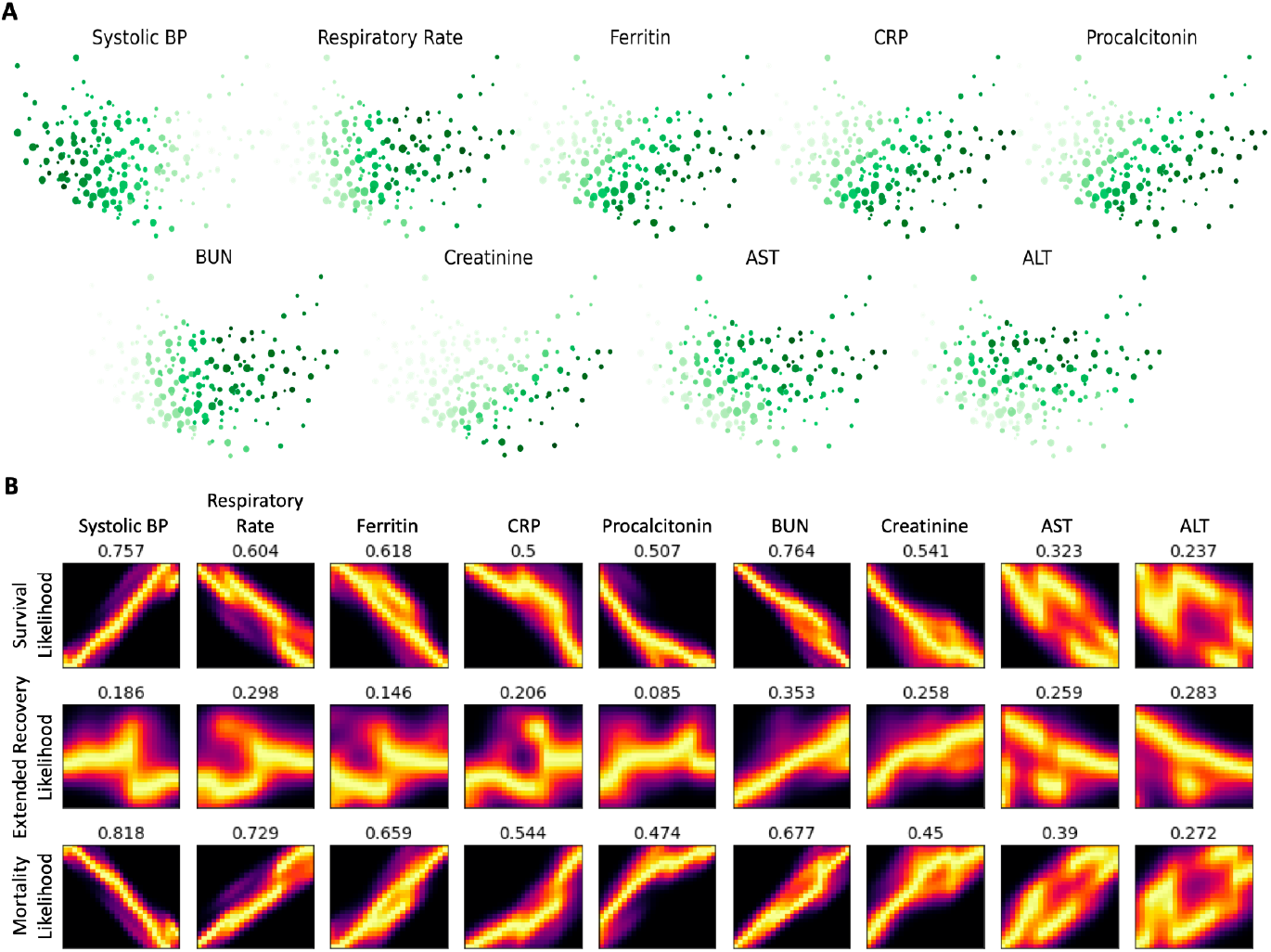
Visualization of multiscale clinical manifold and correlation with patient clinical features. A. Visualizing clinical trends on clinical manifold. B. DREMI and DREVI association analysis between clinical features and patient hospitalization outcome.

